# A hierarchical Bayesian brain parcellation framework for fusion of functional imaging datasets

**DOI:** 10.1101/2023.05.24.542121

**Authors:** Da Zhi, Ladan Shahshahani, Caroline Nettekoven, Ana Lúısa Pinho, Danilo Bzdok, Jörn Diedrichsen

## Abstract

One important barrier in the development of complex models of human brain organization is the lack of a large and comprehensive task-based neuro-imaging dataset. Therefore, current atlases of functional brain organization are mainly based on single and homogeneous resting-state datasets. Here, we propose a hierarchical Bayesian framework that can learn a probabilistically defined brain parcellation across numerous task-based and resting-state datasets, exploiting their combined strengths. The framework is partitioned into a spatial arrangement model that defines the probability of a specific individual brain parcellation, and a set of dataset-specific emission models that defines the probability of the observed data given the individual brain organization. We show that the framework optimally combines information from different datasets to achieve a new population-based atlas of the human cerebellum. Furthermore, we demonstrate that, using only 10 min of individual data, the framework is able to generate individual brain parcellations that outperform group atlases.

## 1. Introduction

The application of machine learning to functional Magnetic Resonance Imaging (fMRI) data promises better models of brain organization. Brain parcellations, which subdivide the brain into a discrete set of functionally distinct regions, are one important type of model with many practical applications. A number of such parcellation schemes have been derived from large resting-state fMRI datasets (Yeo et al., 2011, Buckner et al., 2011, Power et al., 2011, Schaefer et al., 2018, Ji et al., 2019). Previous studies have shown that functional boundaries detected during resting-state are indeed predictive of functional boundaries during task performance (Cole et al., 2014, Laumann et al., 2015, Tavor et al., 2016). However, there is also increasing evidence for systematic differences in the functional organization measured during the task and rest setting (Hasson et al., 2009, Cole et al., 2014, Greene et al., 2020). It is therefore important to consider task-based datasets in deriving brain parcellations (King et al., 2019), foreshadowing a comprehensive understanding of the dynamic nature of the brain’s functional organization.

In recent years, an increasing number of high-quality task-based fMRI datasets that sample a broad range of tasks have become available (King et al., 2019, Nakai and Nishimoto, 2020, Pinho et al., 2018, 2020). Nonetheless, compared to the large and homogeneous restingstate datasets (Van Essen et al., 2013), task-based datasets usually only contain a small to medium number of individuals and are always limited in the tasks that they cover. It would be therefore highly desirable to have a principled way of combining evidence across many datasets into a single model. This is especially important as functional brain organization may not only differ between task and rest, but also between different task sets.

A second important practical problem is that functional brain organization shows considerable inter-individual variations even after anatomical variability is accounted for (Mueller et al., 2013), limiting the usefulness of functional group atlases. This problem could be potentially addressed by including individual resting-state (Wang et al., 2015) or task-based data (King et al., 2019, Pinho et al., 2018, 2020) as a functional localizer to derive individual brain parcellation maps. But a reliable characterization of brain organization requires an extensive amount of individual functional data (Marek et al., 2018), which in practice is often too costly to acquire.

In this paper, we addressed both of these problems by developing a hierarchical Bayesian parcellation framework (Fig. 1), which could be efficiently trained on a range of fMRI datasets, **Y***^s,n^*, recorded in different sessions (*n*) from different subjects (*s*). The model assigns each of the possible brain locations in each individual to one of *K* functional regions (here referred to as parcels). The parcel assignments are collected in the matrices **U***^s^*, with 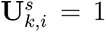 if the *i^th^* brain location is assigned to the *k^th^* parcel. The model estimates the expected value of these latent variables, ⟨**U***^s^*⟩, which provides a probabilistic parcellation for that individual (see Methods 4.1.1 for details). The model consists of a spatial *arrangement model*, *p*(**U**|***θ****_A_*), the probability of how likely a parcel assignment is within the studied population, and a collection of dataset-specific *emission models*, *p*(**Y***^s,n^*|***θ****_En_*), the probability of each observed dataset given the individual brain parcellation. This distributed structure allows the parameters of the model, (***θ****_A_, **θ**_E_*_1_*, ..*) to be estimated using a message-passing algorithm between the different model components (Methods 4.1.4).

**Figure 1:**
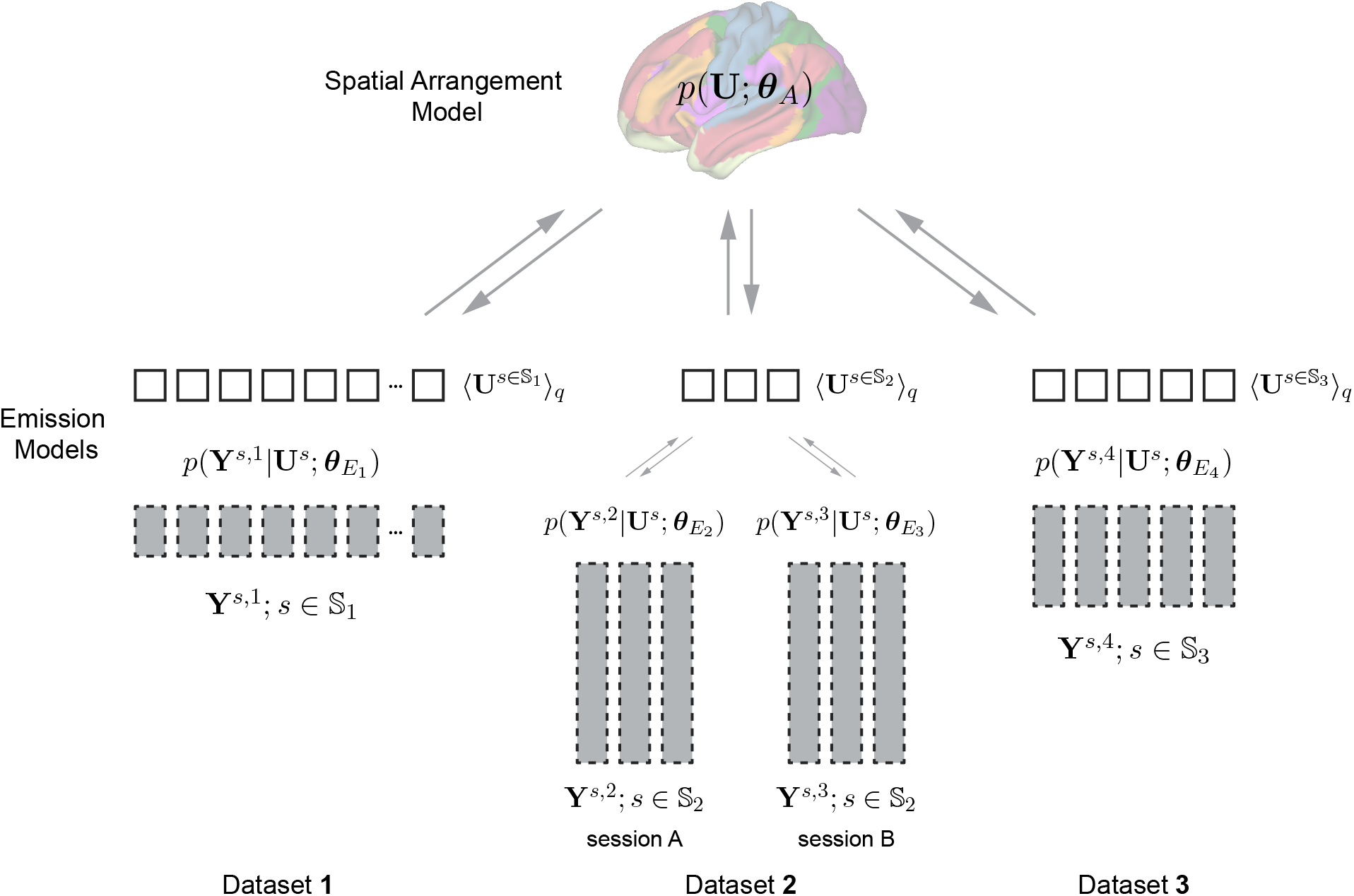
A hierarchical Bayesian parcellation framework for data fusion. Three datasets are shown. Data from each participant are indicated as a gray box. The height of the box indicates the amount of data per participant. Dataset 2 contains two sessions from the same set of participants (*s* ∈ S_2_). The central quantity of the model is the estimated individual brain organization **U^s^**. The spatial arrangement model provides the population-wide probability of observing a specific brain organization.

We applied the new framework to a collection of seven task-based fMRI datasets (Table 1), with four of them containing a wide range of task conditions and three others related to specific functional domains, including executive function and motor movements. Starting within a single dataset, we show that our framework optimally integrates data from a single individual subject with the group-based arrangement model, resulting in substantially improved individual brain parcellations. Then, we compare different approaches to estimate a unified group-based arrangement model across datasets, using both simulations and real data. We show that both group and individual parcellations learned by datasets fusion using our proposed framework outperform the parcellations trained on each single dataset alone.

**Table 1:** FMRI datasets used for the functional fusion. All datasets listed but the last one are task-based, all of them together covering a wide range of psychological domains. The last one refers to resting-state data from a subset of the HCP dataset.

| Name | Subjects | Unique task conditions | Functional scan time (min) | Voxel size (mm) | Description | DOI | Reference |
| --- | --- | --- | --- | --- | --- | --- | --- |
| *MDTB* | 24 | 62 | 240 | 3T, 3mm | Cognitive, motor, perceptual, social | [10.18112/openneuro.ds002105.v1.1.0](https://doi.org/10.18112/openneuro.ds002105.v1.1.0) | King et al. (2019) |
| *Highres-MDTB* | 7 | 9 | 120 | 7T, 1.5mm | Cognitive, motor, perceptual, social | not openly available | not openly available |
| *Nishimoto* | 6 | 103 | 162 | 3T, 2mm | Cognitive, motor, perceptual, social | [10.18112/openneuro.ds002306.v1.0.3](https://doi.org/10.18112/openneuro.ds002306.v1.0.3) | Nakai and Nishimoto (2020) |
| *IBC* | 12 | 208 | 822 | 3T, 1.5mm | Broad Battery | [10.18112/openneuro.ds002685.v1.3.1](https://doi.org/10.18112/openneuro.ds002685.v1.3.1) | Pinho et al. (2018); Pinho et al. (2020) |
| *WMFS* | 16 | 5 + 12 | 5 | 3T, 3mm | Motor and working memory task | not openly available | Shahshahani et al. (2023) |
| *Multi-Demand* | 37 | 12 | 100 | 3T, 2mm | Executive Tasks | not openly available | Assem et al. (2022) |
| *Somatotopic* | 8 | 6 | 96 | 1.8/2.4 | Motor | [10.1152/jn.00165.2022](https://doi.org/10.1152/jn.00165.2022) | Saadon-Grosman et al. (2022) |
| *HCP-Unrelated 100* | 100 | none | 60 | 3T, 2mm | Resting-state | https://www.humanconnectome.org/study/hcp-young-adult/data-releases | (Van Essen et al., 2013) |

## 2. Results

### 2.1. Individual parcellations in the scarce data setting

Given the substantial inter-individual functional variability, it is often desirable to derive parcellations for single subjects. An important limitation, however, is that obtaining a reliable individual parcellation requires a substantial amount of data (Marek et al., 2018). A central feature of our model is that we do not only obtain a parcellation based on the individual data, *p*(**U***^s^*|*Y ^s^*), and a parcellation based on the learned group parameters, *p*(**U***^s^*|***θ****_A_*), but also an optimal integration of individual and the group-level probability map (Methods 4.1.5). We first sought to determine how much improvement this integrated individual parcellation offers. For this purpose, we first trained a group parcellation (17 parcels) on the first task set of the multi-domain task battery dataset (MDTB, (King et al., 2019)). Individual parcellations were derived using between 1-16 imaging runs (10-160 min) of individual training data only. We compared the performance of these ”data-only” parcellations with the group parcellation, and with individual parcellations learned in our framework by Bayesian integration of individual data and group map. All probabilistic parcellations were first converted into hard parcellations using a winner-take-all approach. We then evaluated how well the parcel boundaries corresponded to functional boundaries on the second task set (also 16 runs) acquired on the same subject. For this, we computed the distance-controlled boundary coefficient (DCBC, Zhi et al. (2022), the difference of the within-parcel and the between-parcel correlation of the functional profiles for each spatial distance (see Methods 4.5).

The individual parcellations based on 10 min of imaging data (without using the group probability map, Fig. 2a) performed generally poorly, with an average DCBC of 0.088 (Standard Error of the Mean, SEM = 0.009). Indeed, the individual parcellation performed worse than the group map *t*_23_ = −7.786*, p* = 6.815 × 10*^−^*^8^ (Fig. 2d, dashed line in Fig. 2e). The individual parcellation improved continuously when using more data (Fig. 2b), reaching an average DCBC value of 0.175 (SEM = 0.016) for 160 min of data, ultimately outperforming the group map (*t*_23_ = 3.286*, p* = 0.003). This indicates that there are replicable differences in brain organization across individuals. Individual parcellations can capture these differences, leading to significantly better prediction performance than a group probability map on independent test data.

**Figure 2:**
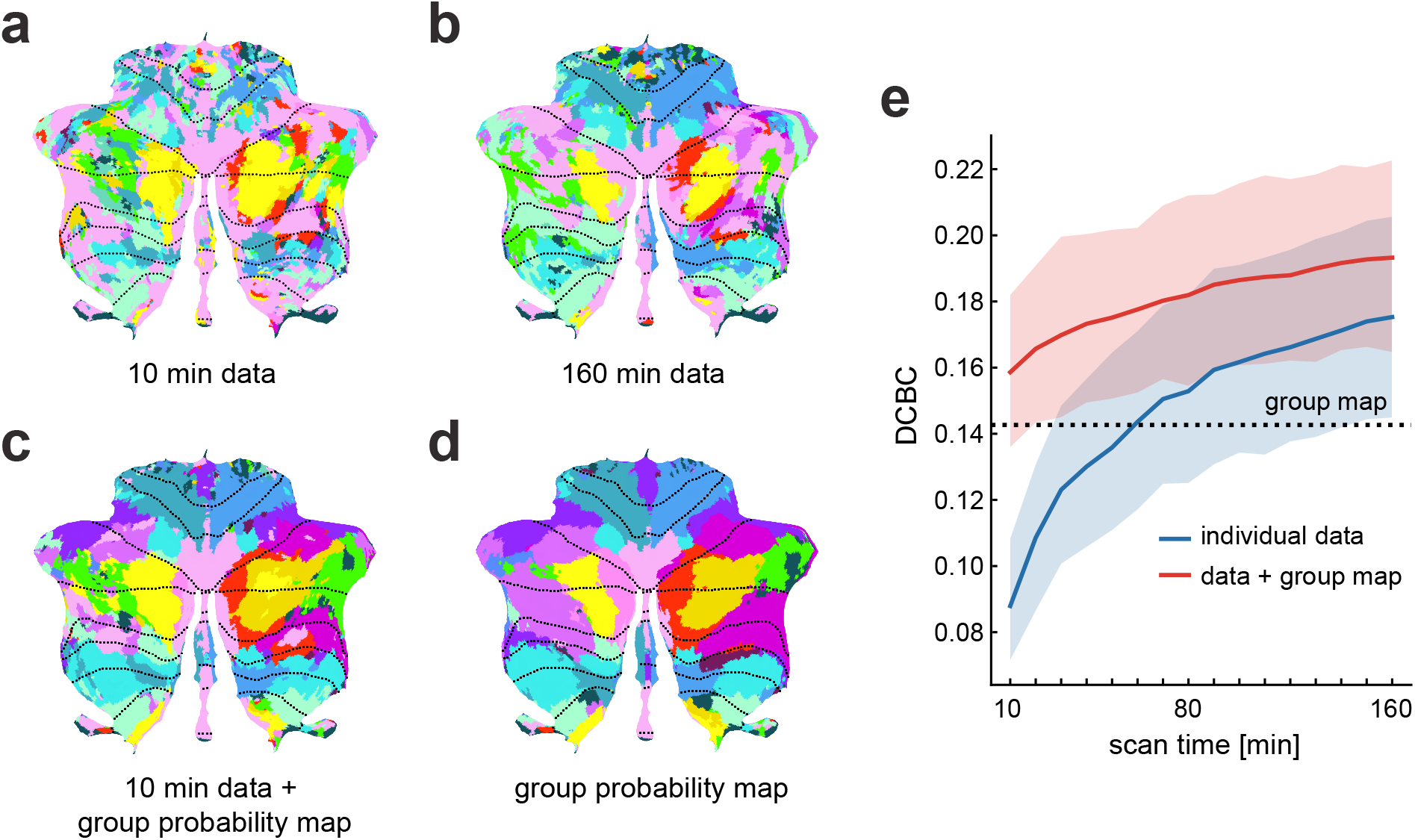
Individual parcellations from the parcellation framework outperform group map. **(a)** An estimated individual parcellation based on 10 min (1 run) of imaging data, using only the individual data. **(b)** An estimated individual parcellation of the same subject based on 160 min (16 runs), using only the individual data. **(c)** The estimated individual parcellation using 10 min of individual data and the group probability map learned by the arrangement model. **(d)** The group probability map from the arrangement model. **(e)** The DCBC value (higher = better) of the parcellations tested on the independent second session of the MDTB data set. Each individual parcellation was estimated using only the individual data (blue curve) or using the individual data and the learned group probability map (red curve). The x-axis indicates the length of the imaging time series (10 min = 1 run) used to estimate the individual parcellations. The error bars represent the standard error of the mean across all 24 subjects.

Although individual parcellations were superior to the group map using more data (blue line in Fig. 2e), our results suggest that more than 110 min of individual imaging data is required to obtain a brain parcellation that is significantly better than the group probability map (*t*_23_ = 2.190*, p* = 0.039). At 60 min of imaging, an individual parcellation map is only just about as predictive as the group probability map. Acquiring this amount of individual data for functional localization is rarely feasible in basic and clinical functional imaging studies.

Our framework, however, automatically integrates the individual data with the group probability map, leading to dramatically improved performance of individual parcellations. For only 10 min of individual data, the DCBC was now significantly higher than the group probability map alone (*t*_23_ = 3.123*, p* = 0.005). Using 10 min of imaging data and our model led to individual parcellation performance that was roughly equivalent to using 100 min of individual imaging data without the model.

The resultant individual parcellation map (Fig. 2c) constitutes an optimal fusion of the individual data and the knowledge learned from the entire group. Even when there was a large amount of individual data available, such as 160 minutes, the integration with the group map led to a significant improvement relative to using only the individual data (*t*_23_ = 5.562*, p* = 1.171×10*^−^*^5^). Another advantage of the integration of group and individual data is that it naturally deals with missing individual brain data. For brain locations (voxels) where the individual data is missing, the group probability map will dictate the parcel assignment.

### 2.2. Dataset-specific emission models optimally capture differences in measurement noise

Different imaging datasets, or even sessions within a single dataset, often show different signal-to-noise ratios. For instance, two different imaging sessions of the IBC data set (Fig. 3a, Methods 4.2) show quite different levels of within-subject reliability, indicative of different levels of measurement noise. A simple approach to modeling different sessions from a single individual is to concatenate the data and model the two sessions with a single emission model (Type 1 model, Fig. 3b). In this scenario, however, it is possible that the second, noisier session will make the integrated model worse than the first session alone. Therefore, in a different version of the model (Type 2), each imaging session was modeled with a separate emission model. This allows differences in variability to be captured by a session-specific concentration parameter (e.g. *κ*^1^ for session 1 and *κ*^2^ for session 2 in Fig. 3c). As long as the *κ*s are estimated accurately, the subsequent Bayesian integration will ensure the optimal weighting across the different sessions. Therefore, even the addition of a low-quality dataset should never lead to decreases in the quality of the integrated model.

**Figure 3:**
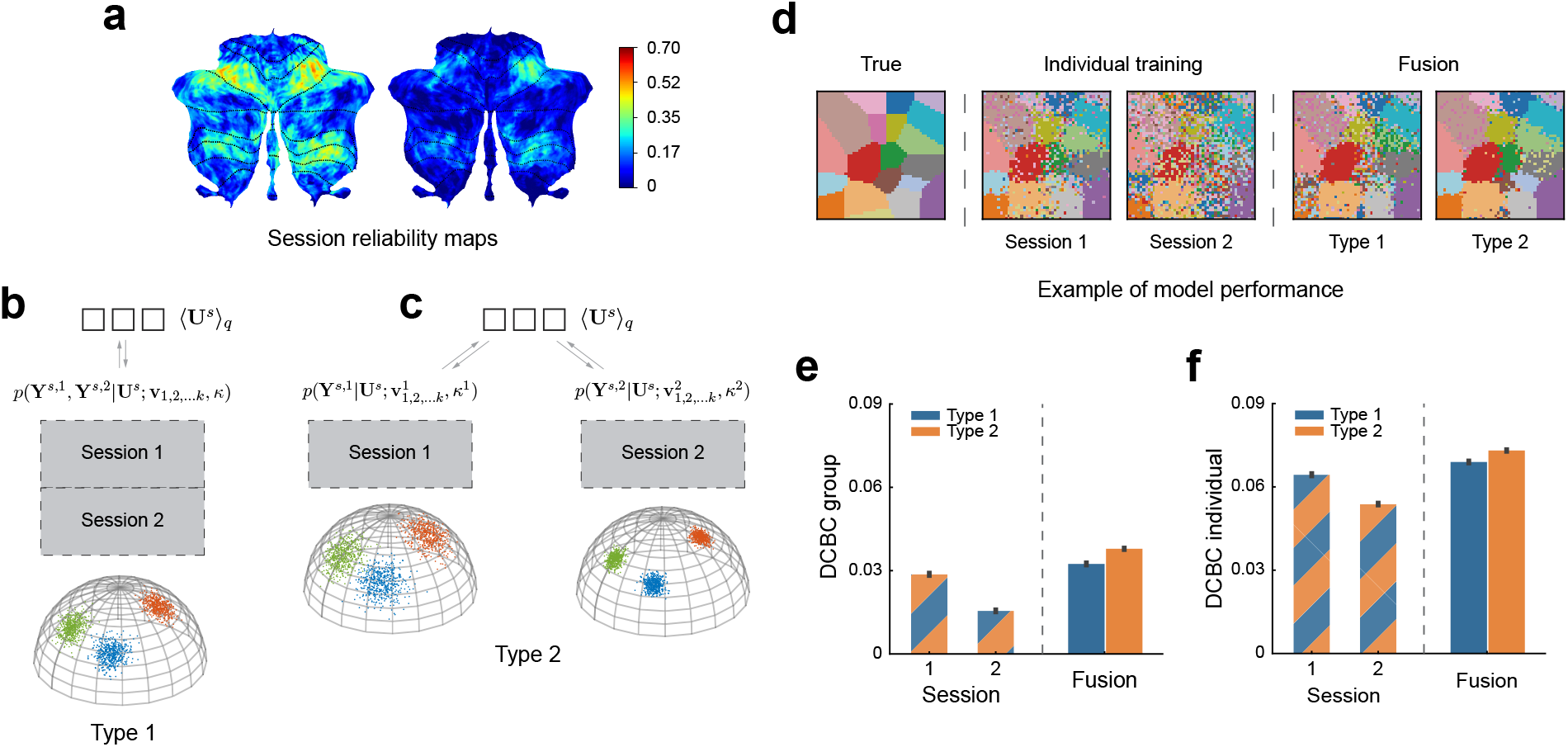
Simulations of data fusion using two synthetic imaging sessions with similar task activation. **(a)** The reliability map of two imaging sessions from the IBC dataset with similar task sets (*hcp1* and *archi*). **(b)** Type 1 model: sessions are concatenated and will be learned in a single emission model. **(c)** Type 2 model: sessions are separated and modeled using two separate emission models. **(d)** Reconstruction of the true parcellation map using synthetic data, using Session 1 or 2 alone vs. the fusion of both sessions using either model type 1 or 2. **(e)** The mean DCBC value of the group map (using model type 1 or type 2) learned from Session 1 or 2 alone or from the fusion of both sessions. **(f)** The mean DCBC value of individual parcellations. Error bars indicate SEM (standard error of the mean) across 100 simulations.

To test for this behavior of the dataset-specific (Type 2) model, we generated two synthetic datasets (sessions) sampled from the same set of subjects with similar task activation but different overall noise variances, 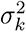 (Methods 4.4). The measurement noise was set to 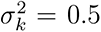 for synthetic session 1 and to 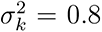 for session 2. We then learned group and individual parcellations using Type 1 or Type 2 models, either using each session alone or fusing both sessions. We tested the performance of all models on an independent simulated test set (Methods 4.4), repeating the simulation 100 times.

The visual inspection of the group parcellations (Fig. 3d) suggests that the group map trained on session 1 alone approximates the true map more accurately than using session 2. The fusion of both sessions improved the group reconstruction, especially when using separate emission models (Type 2). We evaluated the parcellation performances quantitatively using the DCBC measure on the test set (Fig. 3e, 3f). Indeed, both group and individual maps learned from session 1 (DCBC group=0.029, DCBC individual=0.064) showed better performance averaged across 100 simulations than the one using session 2 (DCBC group=0.016, DCBC individual=0.054). When we evaluated the fusion parcellations, the DCBC value of the group and individual map learned by the Type 1 fusion model improved by 0.004 (SD=3.752 × 10*^−^*^3^) and 0.005 (SD=3.781 × 10*^−^*^3^) compared to dataset 1 alone, respectively. The parcellation performance of Type 2 fusion further improves compared to Type 1 by 0.005 (SD=4.006 × 10*^−^*^3^) for the group DCBC and 0.004 (SD=4.666 × 10*^−^*^3^) for the individual DCBC. These simulations demonstrate that session-specific emission models allow for better fusion when the signal-to-noise level differs across sessions or datasets.

### 2.3. Region-specific concentration parameters further improve fusion parcellation

In empirically observed task-based fMRI data, the signal-to-noise level does not only differ between sessions or datasets, but also between different regions within the same session or dataset. Some sessions or datasets provide a better signal-to-noise ratio for some functional regions and a lower signal-to-noise ratio for others. For example (Fig. 4a), the ‘*Preference*’ session of the IBC dataset provided high within-subject reliability in the motor areas, whereas the perspective taking (‘*TOM* ’) session had high reliability in language-related areas. Ideally, a probabilistic framework should account for these differences and optimally combine the region-specific strengths of each dataset. To this end, we introduced a third variant of our emission model (Type 3), which has a separate concentration parameter for each region and session (e.g. 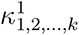 for session 1 and 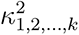 for session 2 in Fig. 4b).

**Figure 4:**
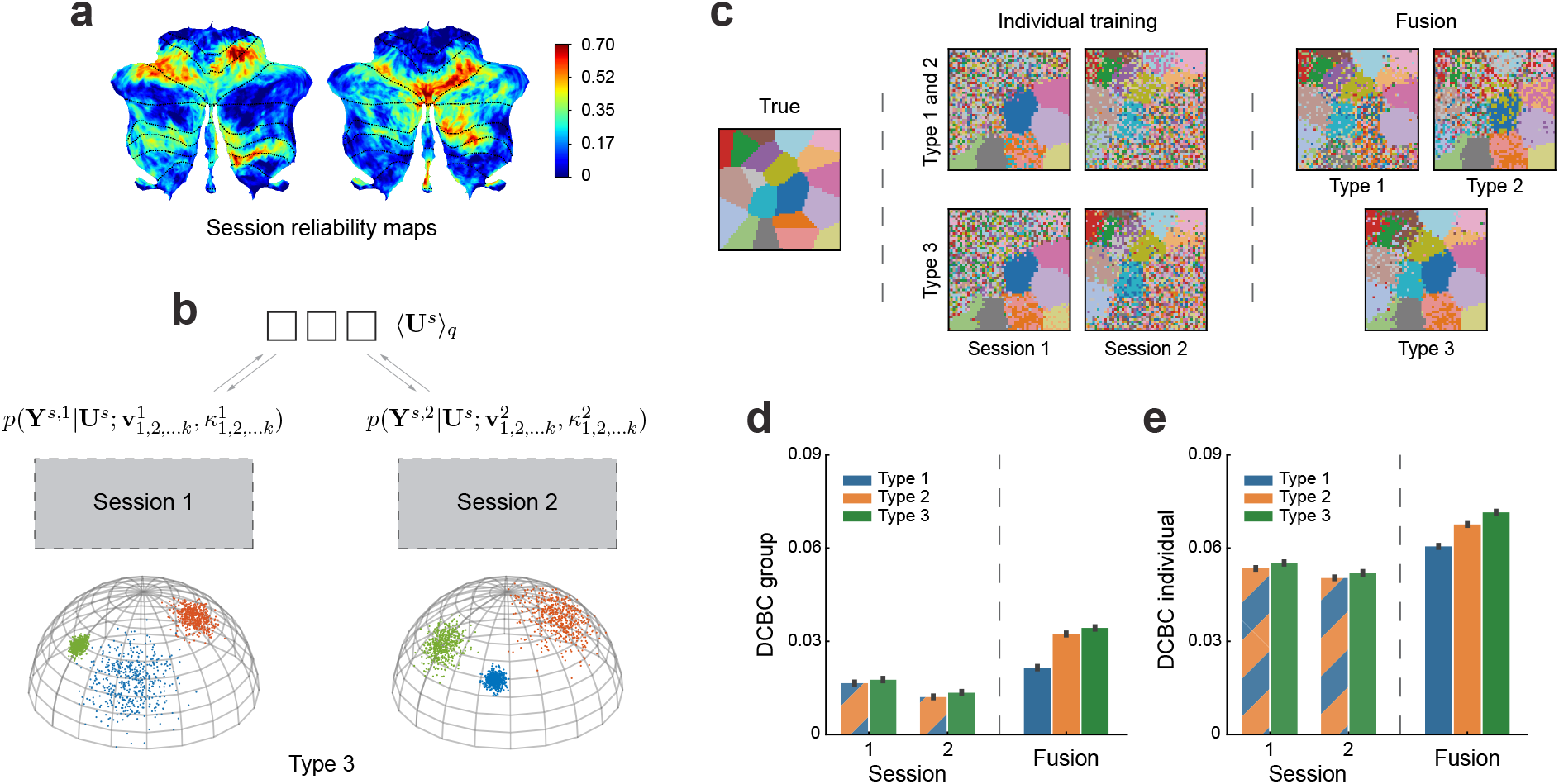
Simulation on two synthetic sessions fusion with different task activation. **(a)** The reliability map of two imaging sessions from the IBC dataset with different tasks (*Preference* and *TOM*). **(b)** Type 3 model: different sessions are modeled using different emission models, and furthermore the concentration parameters *κ*_1,2_*_,…k_* are estimated separately for each parcel. **(c)** The comparison of reconstruction performance when leaned on synthetic session 1 or 2 alone vs. learned by data fusion using type 1, 2, or 3 models. **(e)** The mean DCBC value of the group map across sessions and model types. **(f)** The mean DCBC value of individual maps across sessions and model types. Error bars indicate SEM across 100 times simulation.

To test the ability of this model to pool information across distinct datasets with different types of information, we conducted a second simulation by randomly dividing all functional regions into two groups. Instead of a common signal-to-noise level for all regions, we first created synthetic data in which one session had good signal-to-noise in the first group and poorer signal-to-noise in the other (Methods 4.4). We reversed the assignment for the second synthetic session. When we trained the model on Session 1 or 2 alone, there was high uncertainty of the cluster assignment in the area with low signal-to-noise level (Fig. 4c – *Individual training*). This is no surprise, as the activation here was too weak to detect the boundaries reliably.

Importantly, when combining the two sessions, the functional boundaries that were not detected based on single sessions became visible (Fig. 4c – *Fusion*). However, both Type 1 and Type 2 models needed to compromise: when using session 1 to achieve parcellation of the lower right corner, the same weighting was applied to the upper left regions, decreasing the quality of the parcellation here. In contrast, model Type 3 allowed different weightings in different parcels, using mostly information from session 1 for the lower right parcels and mostly information from session 2 for the upper left regions. The quantitative evaluation (Fig. 4d, 4e) suggests a clear advantage of model Type 3 for both the group (improved 0.002, SD=3.324 × 10*^−^*^3^) and individual parcellation (improved 0.004, SD=3.831 × 10*^−^*^3^).

We also verified the Type 3 model did not perform worse than Type 2 when two sessions had the same signal-to-noise level across all functional regions (see Supplementary Fig. S3). Overall, the model with region-specific concentration parameters showed clear advantages when aggregating across sessions that differ not only in their overall signal-to-noise level, but also in what regions they specifically provide information for.

### 2.4. Model performance on real data and the choice of atlas resolution K

We then attempted to validate the performance of the models on real imaging data. Here, we first used the IBC dataset. This dataset is ideal to test the integration of data from different sessions across the same participants, as it consists of 14 sessions some of which have similar tasks while others do not (Pinho et al., 2018, 2020). We tested the different model types, each time fusing two IBC sessions (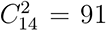 combinations) to learn a probabilistic parcellation model with 17 parcels. The learned models were then evaluated on the six other functional task-based fMRI datasets (see Tabel 1) in terms of their group and individual parcellations. To evaluate the ability of the model to provide individual parcellations, we split each evaluation dataset into two halves. The first half was used to infer the individual parcellations **U***^s^* for the participants of the test set. The other half was used to calculate the DCBC value. We then reversed the role of the two halves and averaged performance across the two cross-validation folds.

We first confirmed that our probabilistic parcellation framework optimally learns group parcellation across sessions when comparing the performance to the parcellation learned from a single session. Specifically, all fusion parcellations showed substantial improvement (Fig. 5) over the best one learned on a single session (Type 1: *t*_98_ = 12.282*, p* = 1.513×10*^−^*^21^, Type 2: *t*_98_ = 18.749*, p* = 3.485 × 10*^−^*^34^, Type 3: *t*_98_ = 15.594*, p* = 2.698 × 10*^−^*^28^). This improvement also held for individual parcellations (Fig. 5b, Type 1: *t*_98_ = 15.283*, p* = 1.100 × 10*^−^*^27^, Type 2: *t*_98_ = 14.198*, p* = 1.624 × 10*^−^*^25^, Type 3: *t*_98_ = 9.353*, p* = 3.079 × 10*^−^*^15^). Additionally, we found the group parcellations learned using session-specific emission models (Type 2) showed significantly better performance than the ones learned by concatenating the data (Type 1) (*t*_98_ = 13.287*, p* = 1.196 × 10*^−^*^23^).

**Figure 5:**
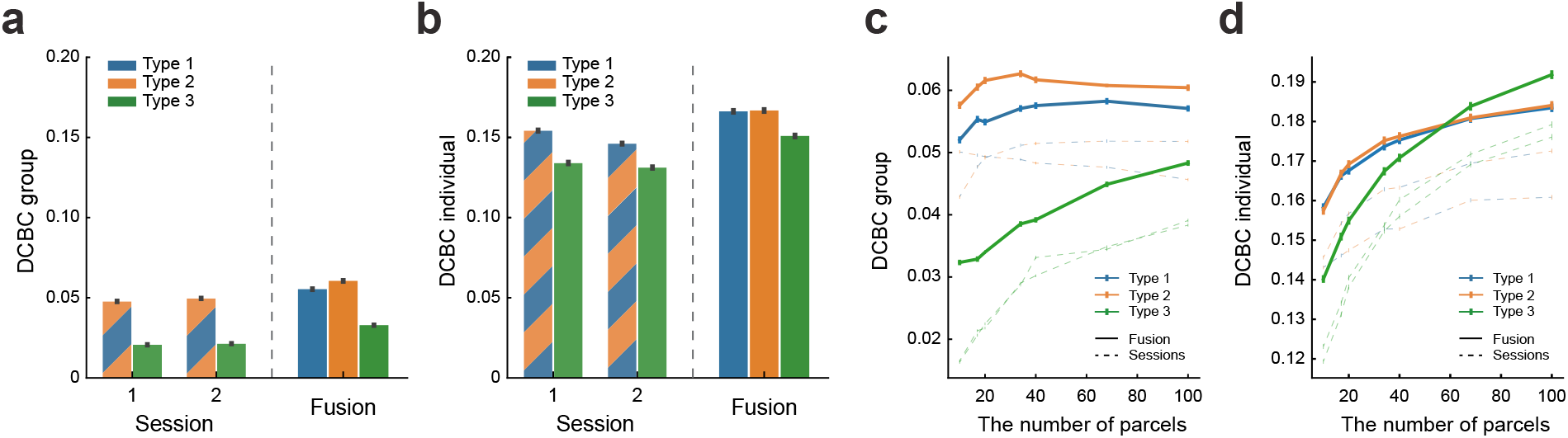
DCBC evaluation of brain parcellations learned on two IBC sessions alone compared to the parcellation learned by fusion. **(a)** Mean DCBC value of the group map across all 91 two-session combinations when training 17-regions parcellations. **(b)** Mean DCBC value of individual maps across all 91 two-session combinations when training 17-regions parcellations. **(c)** Mean DCBC value of the group map as a function of the number of parcels. **(d)** Mean DCBC value of the individual maps across all 91 two-session combinations as a function of the number of parcels. All error bars indicate the SEM across the evaluation subjects across the 6 task-based fMRI datasets.

Against our expectations, however, model Type 3 performed substantially worse on real data when compared to model Type 2 for both group (*t*_98_ = −16.765*, p* = 1.521 × 10*^−^*^30^) and individual (*t*_98_ = −6.269*, p* = 9.807 × 10*^−^*^9^) parcellations. This behavior differed markedly from our simulation results (Fig. 4), where model Type 3 performed consistently better. Further simulations suggested that this behavior can be explained by the choice of the number of parcels (*K*): when *K* was close to or higher than the true number of parcels, model Type 3 outperformed model Type 2. If, however, *K* was chosen to be smaller than the true *K*, model Type 3 started to yield inferior performance (Supplementary Fig. S4). In such cases, one parcel in model Type 3 typically had a very low concentration parameter, effectively capturing all voxels that are unexplained by the model. Model Type 2 constrains all functional regions to have the same concentration parameter, preventing the model from developing a ’residual’ parcel.

This idea suggests that model Type 3 should improve or even outperform model Type 2 when *K* increases and approaches the true number of parcels. Unlike the simulation, the true number of parcels in real data is unknown. We therefore estimated the fusion models on every pair of two IBC sessions using *K* = (10, 17, 20, 34, 40, 68, 100). The evaluation results (Fig. 5c,d) indicated that the performance of the model Type 3 indeed improved with increasing *K*. This improvement was also clearly observed in individual parcellations (Fig. 5d), where the DCBC evaluation of the model Type 3 became as good as model Type 2 around *K* = 60 and showed a significant advantage at *K* = 100 (*t*_98_ = 4.115*, p* = 8.059 × 10*^−^*^5^). A similar pattern exists in the group map evaluation where the averaged DCBC value of 100 parcels substantially improved compared to the ones with only 10 parcels (*t*_98_ = 28.191*, p* = 8.215 × 10*^−^*^49^). For up to 100 parcels, the fusion parcellation from model Type 3 did not appear to be superior to the one from model Type 2 in group evaluation, however, we found this to be the case when considering more datasets (see Fig. 6e).

**Figure 6:**
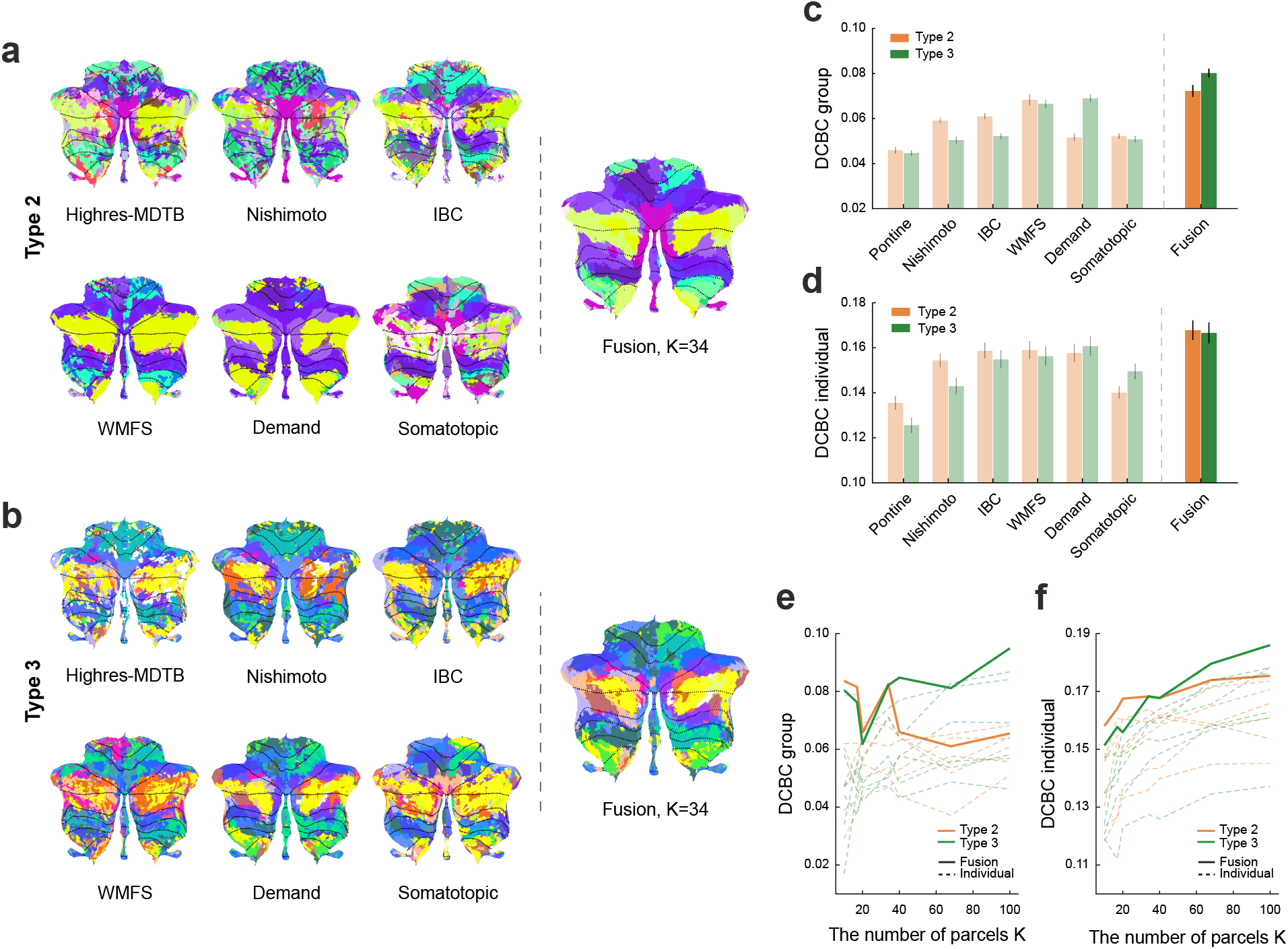
Comparison of cerebellar parcellations learned by type 2 and 3 fusion models using 6 functional task-based datasets. **(a)** The group parcellation maps learned on each individual dataset alone or datasets fusion with 34 parcels using the type 2 model. **(b)** Same as (a), but using the type 3 model. **(c)** Mean DCBC value of the group parcellation maps across subjects in the test dataset. **(d)** Mean DCBC value of the individual parcellation maps across subjects in the test dataset. **(e)** Mean DCBC value of the group map for *K* = 10 to *K* = 100. **(f)** Mean DCBC value of the individual map across for *K* = 10 to *K* = 100.

Overall, across analysis scenarios, we confirm that estimating separate concentration parameters for each session (Type 2) leads to better data fusion on real fMRI data. Additionally allowing a region-specific concentration parameter (Type 3) has both advantages and disadvantages: If the model assumes a large number of parcels, parcellations can improve. If, however, the assumed number of parcels is low, performance appears to be better when constraining the concentration parameter to be the same across regions.

### 2.5. The fusion atlas shows combined strengths across different task-based fMRI datasets

Finally, we trained our fusion model on 6 of the 7 task-based fMRI datasets (Table 1). We reserved the *MDTB* dataset as a test set. The resultant group maps of both models Type 2 and 3 showed the combined strength of the maps trained on individual datasets. For example, only the group map derived from the *Somatotopic* dataset delineated the foot region of the cerebellum (hemispheric lobule IV), while the ones derived from other datasets did not. The Fusion maps (Fig. 6a,b) veridically retained this region. In contrast, the parcellation based on the *Somatotopic* dataset did not show a good parcellation of lobules Crus I and II, but here the fusion map used information from other datasets.

To evaluate the parcellations quantitatively, we calculated the DCBC on the left-out MDTB dataset (Fig. 6c,d). Averaged across all *K*s, all parcellations showed positive DCBC values, which means that the functional boundaries learned from any of the datasets generalized to some degree to the MDTB dataset. The best DCBC among parcellations trained on a single dataset was for the *WMFS* dataset for model Type 2 and for the *Demand* dataset for model Type 3. When we evaluated the fusion parcellations, we found considerable improvements for both models Type 2 and 3 compared to the best individual parcellation. For the fused parcellation using the model Type 2, both the group DCBC (*t*_23_ = 2.339*, p* = 2.840 × 10*^−^*^2^) and the individual DCBC (*t*_23_ = 3.173*, p* = 4.248 × 10*^−^*^3^) were considerably better than for *WMFS*. Similar improvement could be observed for model Type 3, where the fused parcellation significantly outperformed the best single-dataset parcellation (*Demand*) both in terms of the group (*t*_23_ = 7.049*, p* = 3.503 × 10*^−^*^7^) and individual (*t*_23_ = 3.219*, p* = 3.800 × 10*^−^*^3^) DCBC value.

Finally, we compared the fusion across the six task-based fMRI datasets directly between models Type 2 and 3. For *K* = 10, both averaged group and individual DCBC (Fig. 6e,f) were higher for model Type 2 than for model Type 3 (group: *t*_23_ = 0.726*, p* = 0.475; individual: *t*_23_ = 1.842*, p* = 0.078). But when *K* increased to 100, the fusion parcellation for model Type 3 became substantially better than model Type 2 (group: *t*_23_ = 4.551*, p* = 1.426 × 10*^−^*^4^; individual: *t*_23_ = 2.468*, p* = 2.144 × 10*^−^*^2^). The cross-over occurred somewhere around *K* = 34, where models performed equivalently (group: *t*_23_ = 0.210*, p* = 0.835; individual: *t*_23_ = −0.009*, p* = 0.993).

### 2.6. Integrating resting-state data into the task-based parcellation

Lastly, we investigated the fusion of resting-state and task-based data into a single parcellation atlas. To do so, we used the cortical connectivity profile for each cerebellar voxel derived for 50 participants from the HCP*Unrelated 100* dataset (see Method 4.2). As we wanted to evaluate performance on a large range of task-based datasets, we trained the model on 6 out of 7 task datasets and evaluated the performance on the left-out task dataset. We then repeated this scheme for all 7 task datasets. The combined and resting-state parcellations were also trained and evaluated in a similar approach (for the resting-state parcellation, no dataset had to be left out).

Averaging the DCBC evaluations across models Type 2 and 3, the models trained on the combination of resting-state and task-based datasets outperformed the ones trained on resting-state or task-based datasets alone. For the group parcellation (Fig. 7a), the combined model was significantly better than the one trained on resting-state (*t*_110_ = 6.349*, p* = 4.983× 10*^−^*^9^), and task-based datasets (*t*_110_ = 3.886*, p* = 1.745 × 10*^−^*^4^). Similar results were found for individual parcellations (Fig. 7b, vs. resting state alone: *t*_110_ = 7.625*, p* = 9.287 × 10*^−^*^12^, vs. task-based alone *t*_110_ = 7.254*, p* = 6.027 × 10*^−^*^11^.

**Figure 7:**
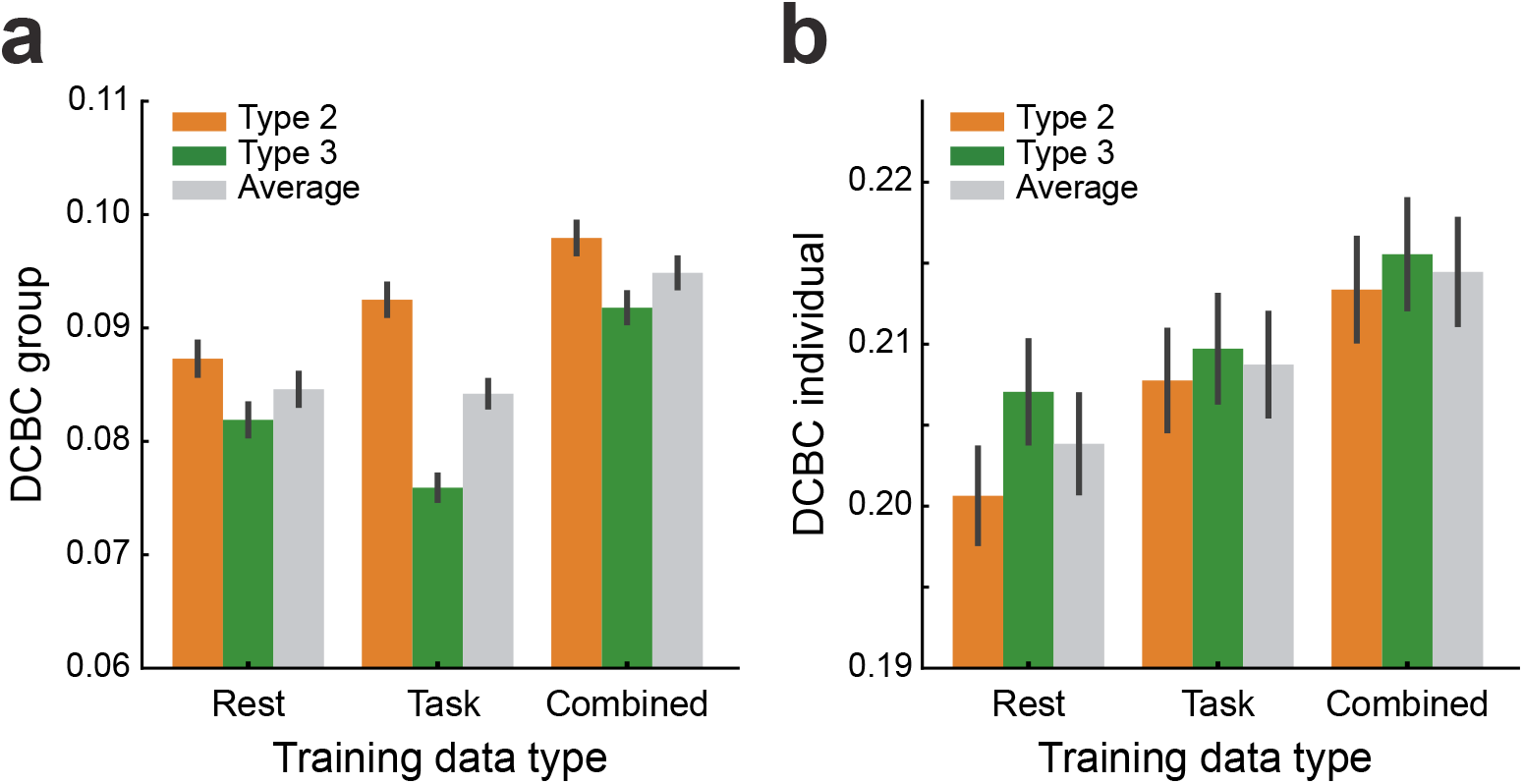
Performance comparison of the cerebellar parcellations using resting-state data only, task-based data only, or the combination of both. Probabilistic parcellations were learned using Type 2 (orange) or 3 (green) models. The gray bar indicates the averaged performance across the two models. **(a)** Mean group DCBC, and **(b)** mean individual DCBC evaluated on the task-based datasets in a leave-one-dataset out fashion. Error bar indicates the SEM across all 111 subjects of the 7 task datasets

## 3. Discussion

We developed a hierarchical Bayesian framework to learn probabilistic brain parcellation by fusing data from both functional task-based and resting-state fMRI datasets. Our work introduces two main innovations: First, by dividing the problem into a common spatial arrangement model and a set of dataset-specific emission models, we are able to optimally integrate information across many, quite heterogeneous, datasets. Second, because the framework directly models individual differences in brain organization, it provides not only a probabilistic group atlas, but also allows the user to obtain an optimal estimate of brain organization for new individuals.

### Learning functional brain parcellations across datasets

While most of the current brain parcellations are generated using functional resting-state fMRI data, a number of studies (King et al., 2019, Cole et al., 2014) suggest that boundaries derived using restingstate data can differ systematically from those measured during task performance. One possible interpretation of this finding is that the boundaries of functional regions truly shift depending on the task the person performs (Salehi et al., 2020). However, given that there is a basic common organization that is stable across rest and different tasks (King et al., 2019, Tavor et al., 2016), an alternative interpretation is that the boundaries stay the same, but are more or less visible depending on the task sets or mental states (such as rest) during which they are measured. This is obviously true for task sets that emphasize one specific aspect of mental function (See Fig. 4a), but also applies to resting-state data. For example, in resting-state data, the left and right-hand regions are usually highly correlated and often end up in the same parcel. However, when using a task set that contains both left and right unimanual movements, the two regions are readily dissociated (King et al., 2019). Therefore, the integration of data from a large array of tasks promises a more representative map of brain organization.

Because there is no single, large task-based dataset that would cover all of the mental functions, we developed here a framework that allows us to fuse data from a growing number of deep-phenotyping task-based datasets with fewer participants (King et al., 2019, Pinho et al., 2018, 2020, Nakai and Nishimoto, 2020, Assem et al., 2022). To make data fusion feasible in a Bayesian framework, we deployed a series of emission models, each one learns the specific characteristics of the corresponding dataset, including the expected response for each brain region and their variability. The integration across datasets is achieved through a common spatial arrangement model, which characterizes the variability of the functional organization across individuals. As shown in the simulations (Results 2.2 and 2.3), this allows us to integrate the strength of different datasets without inheriting their weaknesses. We can now deploy this framework to an increasing number of real datasets, namely the “wide” datasets with many participants (King et al., 2019), and “deep” datasets with only a few participants but a detailed characterization of each studied individual (Nakai and Nishimoto, 2020, Pinho et al., 2018, 2020). Given the message-passing algorithm (Methods 4.1.4), the individual datasets do not necessarily need to be hosted on the same server, but each dataset and emission model can be housed separately. This architecture will promote the scaling of the approach as it allows for distributed computing across many sites, which will be necessary to finally approach a “big-data” regime for learning complex models of functional brain organization. The distributed nature of data storage also makes the framework more suitable for clinical data, which for data privacy reasons often needs to remain within a dedicated server infrastructure.

### Individual vs. group parcellation maps

Group parcellation maps identify patterns of functional organizations that are common and consistent across individuals. Group parcellations are in common use, as they provide a consistent framework to analyze and report functional imaging data, and can be applied using only the anatomical image from the individual. However, the boundaries between functional regions vary substantially across individual brains (Braga and Buckner, 2017, Gordon et al., 2017, Kong et al., 2021), possibly biasing subsequent analysis (Bijsterbosch et al., 2018, 2019). Recent studies suggest that the inter-individual difference may be even more pronounced in the human cerebellum (Marek et al., 2018). Therefore, using individual brain parcellations has the potential to improve the precision and quality of subsequent analyses. A major limitation, however, is that a substantial amount of individual data is necessary to derive an individualized map of sufficient quality (Marek et al., 2018). In our study, we found that 60 minutes of individual data were required to reach the same performance as the group map, and more than 110 minutes were necessary to substantially outperform it (see Results 2.1). For most studies, acquiring this amount of data for an individual functional localizer would be prohibitive, explaining the persistent popularity of group maps.

Different from previous approaches to leverage the group and individual parcellations (Salehi et al., 2018, Zhang et al., 2021), our approach performs the combination in a principled (Bayesian) way, weighting each part according to the respective uncertainty. Even when using a very short functional localizer (10min), the resultant individual parcellation outperforms the group map. Relative to parcellation built on individual data only, we found that the integrated estimate had a performance equivalent to using 100 min. Finally, the Bayesian approach also automatically deals with missing data from individuals due to lack of coverage or signal dropout.

While we derived and tested the individual parcellations for participants included in our training set, the model can also be used to derive individual parcellations on completely new participants. This would only require researchers to estimate a new emission model for the specific task set used for the functional localization data. In this process, the parameters of the arrangement model, which was trained across datasets, can be frozen. Therefore, an efficient estimation can be achieved even for small groups of participants, and results can be interpreted in the framework of established atlases. This approach makes individual functional localization in larger studies feasible, which is important for practical and clinical applications.

### Comparing dataset-specific and regions-specific uncertainty parameters

The concentration parameter (*κ*) in each emission model dictates how strongly the respective dataset is weighted, both when learning to determine the group parcellation map, and when deriving an individual parcellation. In this paper, we tested three ways of estimating this concentration parameter: (a) we simply concatenated all sessions for each subject, giving the entire dataset a single concentration parameter (Type 1); (b) we used a separate emission model and, therefore, a separate concentration parameter for each session (Type 2); and (c) we used a separate concentration parameter for each session and region (Type 3).

We first showed that the model Type 2 performed better than model Type 1 in capturing different levels of measurement noise from different sessions in both simulation and real data (Results 2.2, 2.4). However, when we compared Type 2 (dataset-specific) and Type 3 (region-specific) models, we found that each had specific advantages, which is also dependent on the choice *K*, the number of parcels (Results 2.4). When allowing separate concentration parameters for each session and region (Type 3), we can account for the fact that some sessions may contain tasks that provide signals in some areas, while other sessions may highlight other areas. This example clearly the case in the IBC dataset (Fig. 3a). While in simulation, model Type 3 led to superior performance, on real data it often performed worse than model Type 2. In model Type 3, we found that when the assumed number of parcels (*K*) was smaller than the true number of parcels, one region would be estimated to have a very low concentration parameter, such that it could model all the residual, non-explained regions. Such a residual region led to a more fragmented group parcellation (Fig. 6b) and an impaired evaluation of the independent data.

However, the constraint of equal concentration parameters across all regions (model Type 2) prevented this from happening. This led to compact clusters regardless of the choice of *K* (Fig. 6a). Nonetheless, for large *K*, model Type 3 could outperform model Type 2. The choice of emission model (Type 2 or Type 3) therefore will depend on desired granularity of the parcellation and likely also the amount and quality of data available. Our framework offers both implementations, allowing the user to choose the correct algorithm in a context-specific manner.

### Choice of datasets: task-based vs. resting-state fMRI

Our evaluation of task-based and resting-state parcellations (Fig. 6) shows that both can predict the functional boundaries measured in a left-out task set well above chance. A visual inspection of the two parcellations (Supplementary Fig. S5a,b), however, also reveals some systematic differences (King et al., 2019, Cole et al., 2014). One important decision when applying our framework is therefore which datasets to include. Our Bayesian framework weights each dataset according to its reliability. Because different datasets will emphasize different sets of functional boundaries, and because the true number of functional regions is likely larger than the number of assumed parcels, each dataset will bias the final parcellation in a specific direction. A single large dataset could dominate the group map, possibly reducing the predictive performance for other datasets. It is therefore important to achieve a good balance between resting-state and task-based datasets highlighting different cognitive domains (Salehi et al., 2020). Where this balance lies, or whether it is preferable to have different brain parcellations for different functional states, remains a research question that demands further attention.

### Limitations and further developments

Being able to leverage an increasing number of datasets will hopefully also allow the development of models that can learn regularities in the spatial arrangement of functional regions in the human brain. In this paper, we have used only the independent spatial arrangement model, which in essence learns a probabilistic group atlas. In our framework, however, we can also use models that make assumptions about the intrinsic smoothness of individual functional parcellations, such as a Markov Random Field (MRF) spatial prior (Ryali et al., 2013, Schaefer et al., 2018, Kong et al., 2019) with coupling parameters. As a further extension, deep generative models, such as a deep Boltzmann machine (Salakhutdinov and Hinton, 2009), provide a promising avenue to actually learn the complex short- and long-range dependencies in functional brain organization. While the emission models would remain the same, both the E-step and M-step for the spatial arrangement model would rely on an approximation through sampling. Training such models will require a large amount of data, and our framework takes a critical step in this direction by enabling the integration of a wide range of varied datasets.

Another direction for possible improvements is to explore other forms of emission models. Here, we used a mixture of vMF distributions (Methods 4.1.2), which for both resting-state and task-based data has been shown to perform considerably better than a mixture of multivariate Gaussians (Røge et al. (2017), Supplementary Fig. S1). In contrast to resting-state data, task-based data provide not only a direction of the functional profile, but also a signal amplitude, as the experimental paradigm allows for separate estimation of signal and noise. The signal amplitude could be potentially used to distinguish between reliable and noisy brain locations: we found that the functional profile at brain locations with larger signal magnitudes tended to be closer to the mean response vector for that region. Nonetheless, the vMF model ignores this information as each profile is standardized to unit length. During the development of the model, we therefore experimented with weighted vMF models, in which the emission log-likelihood from each brain location was weighted by its respective signal-to-noise level. In practice, however, we found the final performance of the model did not improve enough to outweigh the possible instabilities in the estimation of the weights. We then decided to stay with the standard vMF mixture model for this paper. But, an emission model with voxel-, region-, and subject-specific signal-to-noise parameters might be useful for certain applications.

### Conclusion

This article designs and evaluates a hierarchical Bayesian parcellation framework for data fusion across heterogeneous data sources. In conjunction with a collection of task-based and resting-state datasets which were preprocessed and stored in a consistent manner, the framework enables optimal estimation of functional brain organization across a range of diverse datasets. Here, we have applied the framework to derive new functional maps of the human cerebellum - however, the same process could be repeated nearly effortlessly for any other brain structure.

We anticipate that this framework will be useful for two reasons. First, the model can provide individual functional parcellations for new subjects using very limited individual data. While normally individual parcellations require an extensive amount of data (Marek et al., 2018), our framework makes it feasible to derive an individual region definition of considerably better quality than a group map with 10 min of functional localizer data. Secondly, the framework allows the optimal fusion of functional insights using a range of different task-based datasets, thereby overcoming the limitation that current task-based datasets are restricted both in terms of the breadth of their task battery and the number of subjects. The framework accurately quantifies the different signal-to-noise levels across sessions and datasets, thereby providing an optimal weighting for each. The resultant maps possess a combined strength in detecting the detailed functional boundaries, outperforming the parcellations trained by single datasets.

## 4. Methods

### 4.1. A hierarchical Bayesian parcellation framework for data fusion

We introduce here a hierarchical Bayesian framework that can be used to learn a probabilistic brain parcellation across multiple fMRI datasets. The framework (Fig. 1) consists of a group-based brain parcellation model (the spatial arrangement model), and a series of dataset-specific emission models. The two parts of the framework are connected by a message-passing and collaborative-learning process, making learning and inference computationally efficient.

The framework is able to learn parcellations from a collection of data **Y***^s,n^* recorded from different subjects (*s*) during different experiments or sessions (*n*). S*_n_* is the set of subjects for the *n*-th experiment or session, and S := {S_1_ ∪S_2_ ∪*…*∪S*_n_*} is the entire set of unique subjects. The parcellation model assigns each of the *P* possible brain locations in each individual *s* to one of *K* functional regions (here referred to as parcels). The parcel assignment for the *i*-th brain location is denoted in the one-hot encoded vector 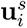, and collected into the *K* × *P* matrix **U***^s^*. This individual brain organization is the central latent variable in the model. The model estimates the expected value, ⟨**U***^s^*⟩, which provides a probabilistic parcellation for that individual - specifically 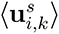 is the probability that brain location *i* is part of the functional region *k*. Note that we use ⟨·⟩ to denote the expected value throughout.

The arrangement model provides a probabilistic group model of how likely a certain parcel assignment to brain locations is across individuals, *p*(**U**; ***θ****_A_*). This probability depends on a set of (to-be-estimated) parameters of arrangement model (***θ****_A_*). In this paper, we use a spatial arrangement model that estimates these probabilities for each brain location independently (Methods 4.1.3), and therefore effectively learns a group-based probabilistic brain atlas (see discussion for further extensions).

Each emission model specifies the likelihood of observed data given the individual brain parcellation, *p*(**Y***^s,n^*|**U***^s^*; ***θ****_E_*). For each dataset/session, we introduce a separate emission model with a separate set of emission-model parameters (***θ****_E_*). This allows us to model the different task sets with different signal-to-noise levels inherent in each experiment/session.

#### 4.1.1. EM algorithm for Probabilistic parcellation with data fusion

We used an *Expectation Maximization* (EM) algorithm to optimize the parameters (***θ***) of the hierarchical Bayesian model. For such models, the computation of the log-likelihood of the data, log *p*(**Y***^s^*; ***θ***), is unfeasible given a large number of latent variables in the model (here, these are the individual brain parcellations **U***^s^*).

The key idea in EM is to introduce a proposal distribution over the latent variables *q*(**U**), and then to optimize the *Evidence Lower Bound* (ELBO) of the model (Wainwright et al., 2008, Blei et al., 2017). The ELBO provides a lower bound to the full likelihood (over all datasets and subjects) that we want to optimize:

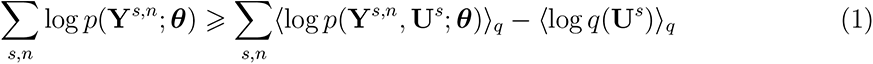

The first term of the ELBO is the expected complete log-likelihood L. Given the model structure, this quantity can be further split into the expected emission log-likelihoods L*_En_* for each experiment or session and the expected arrangement log-likelihood L*_A_* as:

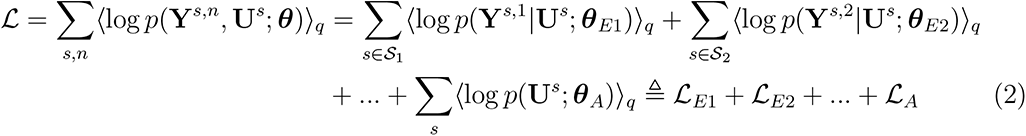

where the parameters are subdivided into those for the arrangement model, ***θ****_A_*, and those for each of the emission models {***θ****_E_*_1_*, **θ**_E_*_2_*, …*}. This division makes the parameter updates that can be performed independently for the arrangement and emission models.

In the expectation step, the ELBO is increased by updating the proposal distribution *q*(**U***^s^*) to the approximate posterior distribution, given the current set of parameters as

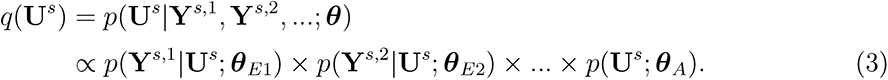

This step also allows us to calculate the expectation of the latent variables, resulting in an estimate of the individual brain parcellations ⟨**U***^s^*⟩*_q_*. In the maximization step, we update these parameters using these estimated individual brain parcellations.

#### 4.1.2. Dataset-specific emission models

One commonly-used choice for a model of fMRI data across regions is the *Gaussian Mixture Model* (GMM) (Golland et al., 2008). However, the amplitude of fMRI brain signals **y***_i_* (whether or not they are normalized by the measurement noise) vary greatly between datasets, participants, and brain locations. That is, two voxels in the same region may have highly correlated signals, but the amplitude of one may be twice as large as the other. Therefore, an increasing number of modeling approaches for resting-state fMRI data use a mixture of *von Mises-Fisher* (vMF) distributions (Banerjee et al., 2005, Ryali et al., 2013, Schaefer et al., 2018, Lashkari et al., 2010, Yeo et al., 2011). It has been demonstrated that such a directional distribution outperforms the GMM in modeling resting-state fMRI data (Røge et al., 2017). Here, we confirmed this is also the case for task-based fMRI data: the vMF mixture model performed better in model evaluation than the GMM (Supplementary Fig. S1). We thus used the vMF mixture as our primary emission model.

The probability density function of a *N* -dimensional (*N* ⩾ 2) vMF distribution for a data point **y***_i_*(∥**y***_i_*∥ = 1) is defined as:

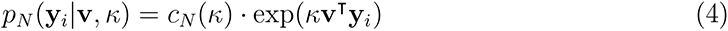

where **v** denotes the mean direction (∥**v**∥ = 1), *κ* indicates the concentration parameter (*κ* ⩾ 0). The normalizing constant *c_N_* (*κ*) is given by:

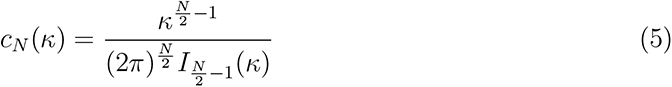

where *I_r_*(·) refers to the modified Bessel function of the *r* order. In a *k*-classes vMF mixture, each of the 1 ⩽ *k* ⩽ *K* parcels is specified with the parameters {**v***_k_, κ_k_*}, where *κ_k_* is the concentration parameter and **v***_k_* is the mean direction vector. Because any spatial dependency in the data is modeled through the arrangement model, these emission log-likelihoods can be computed separately for each brain location *i*. For each subject *s* and emission model *n*, we can calculate the data log-likelihood for each *i* brain location as:

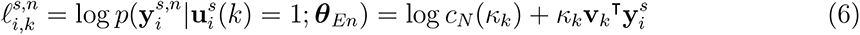

We explored three variants of this model: (a) **Type 1** model assumes that all sessions of a single subject are concatenated and modeled with a single emission model; (b) **Type 2** model uses different emission models for different sessions from the same participant (Fig. 1, Dataset 2). Evidence from different sessions of the same subject is combined during the message passing (eq. 3). Different sessions have different concentration parameters *κ*, providing the possibility of adaptive weighting across sessions. The concentration parameter, however, is assumed to be the same across all parcels; (c) **Type 3** model is identical to the Type 2 model, but employs a different concentration parameter, *κ_k_*, for each of the parcels. In the maximization step, the emission model parameters ***θ****_E_* := {**v***_k_, κ_k_*} are updated by maximizing the expected emission log-likelihood L*_E_* (Supplementary materials S2).

#### 4.1.3. The spatial arrangement model

The arrangement model aims to provide a (possibly not normalized, see discussion) probability measure *p*(**U**; ***θ****_A_*) for each unique individual *s* (*s* ∈ S) in the studied population over a set of latent variables **u***^s^*, which indicates the affiliation of a certain brain location *i* to a specific functional region *k*. We considered here the most basic architecture for the spatial arrangement model, namely the **independent arrangement model**, where different brain locations are considered to be mutually independent. In this case, the spatial arrangement model learns the group probability at each location *i* across all subjects belonging to a parcel *k*, denoted as *p*(**u***_i_*(*k*)). We parameterize this model using a group log-probability parameter *η_i,k_* for each brain location *i* and parcel *k*:

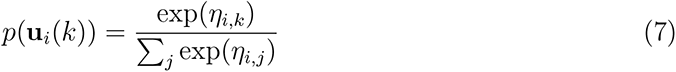

This arrangement model can be estimated using the EM algorithm for inference. In the Estep, we calculate the posterior 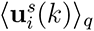 for each individual by integrating the log evidence from the data and the group prior parameter *η_i,k_*:

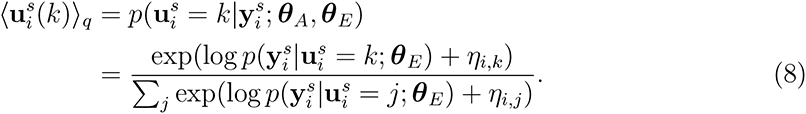

The arrangement model parameters ***θ****_A_* := {*η_i,k_*} are then updated in the M-step (Supplementary materials S6).

#### 4.1.4. Message passing and collaborative learning

Since the full model breaks into different parts (Fig. 1), the learning algorithm can be partitioned into separate E-steps and M-steps for arrangement and emission models (Algorithm 1). The two parts then communicate through a message-passing process. Specifically, if there are multiple emission models in the framework, each of the *n* emission models calculates 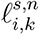 (eq. 6) for each individual *s*.

During *message-passing*, the evidence for a single subject *s* is integrated across any experiment/session that is available for this subject (e.g. Dataset 2 in Fig. 1),

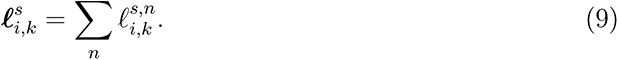

These combined emission log-likelihoods 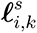 are then collected and passed to the arrangement model. The arrangement model then computes the posterior expectation 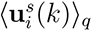 (eq. 8) of the parcel assignment in each subject *s*, which are collected into a *K* × *P* matrix ⟨**U***^s^*⟩*_q_*. These quantities are then used to calculate the expected emission log-likelihoods L*_En_* and the expected arrangement log-likelihood L*_A_*. In case of an independent arrangement model, the expected arrangement log-likelihood L*_A_* can be computed in closed form:

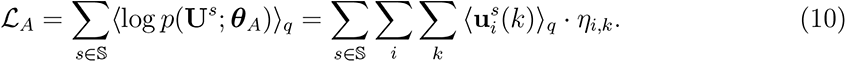

Similarly, the expected emission log-likelihood is calculated by multiplying the data loglikelihood in eq. 6 with the posterior expectation (eq. 8) and summing these quantities over subjects, brain locations, and parcels:

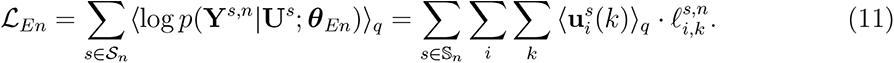

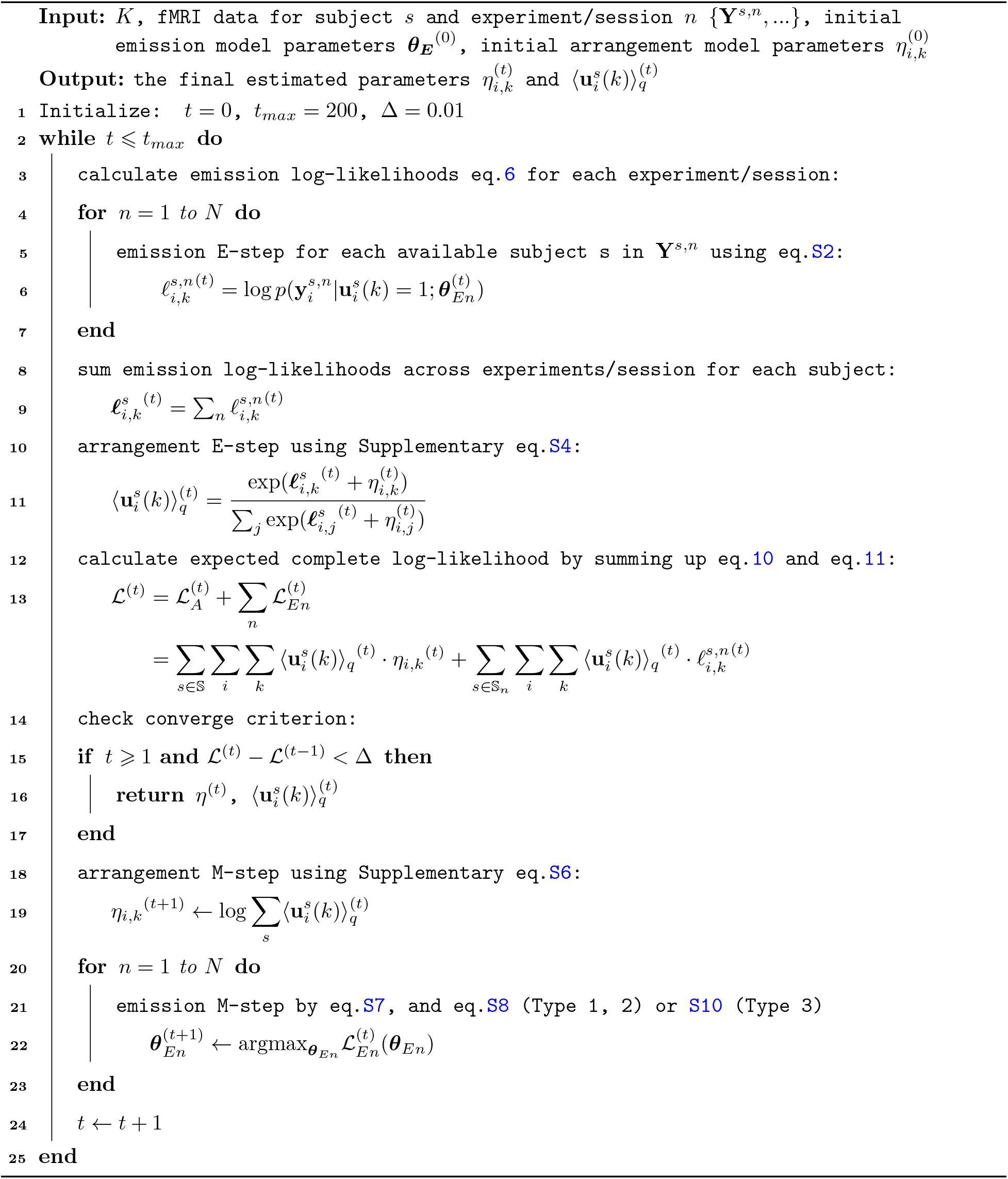
Algorithm 1: EM algorithm of the fusion framework.

The sum of these expected log-likelihoods L (in eq. 2) is then used as an objective function to check the convergence.

In the implementation, the algorithm takes inputs of the fMRI datasets **Y***^s,n^* (*n* = 1, 2*, …, N* and *s* ∈ S) with the initial arrangement and emission model parameters, 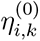 and 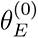. The parameters 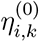 were initiated randomly from a normal distribution. For the initial emission model parameters, the mean direction vectors 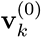 were also drawn from a normal distribution and normalized to be unit vectors. The initial concentration parameters *κ_k_*^(0)^ were randomly drawn from a uniform distribution between 10 to 150, as we wanted to start with a ‘medium-sized’ directional variance. After convergence, the algorithm returns the estimated group parameters for arrangement and emission models, as well as the posterior expectation 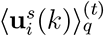 from the last iteration (*t*).

#### 4.1.5. Individual and group parcellations

Once the model is trained, the group probability map can be derived from the estimated parameters of the spatial arrangement model. For the independent arrangement model, the *k*-long vector of probabilities at each brain location is 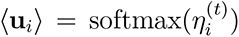. To obtain a hard parcellation for later evaluation (Methods 2.5), we applied a winner-takes-all approach assigning each brain location *i* to the parcel with the highest probability (arg max*_k_*⟨**u***_i_*(*k*)⟩).

Individual parcellations can be obtained even for the individuals that were not part of the model training by applying a single E-step, using the trained parameters and their data. This procedure effectively integrates the individual data likelihood with the group probability map. Since we assume an independent spatial arrangement model, the posterior expectation for a location *i* of subject *s* can be exactly computed as,

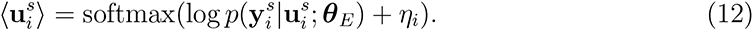

Similarly, a hard individual parcellation was then again obtained by assigning *i* to the region with the highest probability, 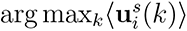. For the comparison reported in section 3.1, we also derived a parcellation only based on data likelihood without taking the group probability into account:

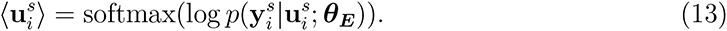

#### 4.1.6. Initialization and convergence

As for most other complex non-convex optimization problems, local minima and slow convergence also constitute a problem during learning in our framework. While each emission model quickly learns a set of mean vectors **v***_k_* that reasonably approximates the respective dataset, the different parcels are not necessarily aligned across the datasets. This is especially the case when the emission models are randomly and independently initialized. As the arrangement model receives conflicting information from different emission models, it can take a long time to bring the different emission models into alignment.

To solve this problem, it is sufficient to start the algorithm with a single down-pass of information from the (randomly initialized) arrangement model to all emission models. That is, during the first iteration of the loop, we skipped the calculation of the emission log-likelihood (line 3-9) of the Algorithm 1, setting all 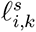 to zeros. This “pre-training” helps to align the corresponding parcel assignments across all datasets.

A further technique to address the slow convergence is to initialize the model from many different random starting points, and only perform a few learning iterations. After this initial phase of learning, we picked the model with the highest expected log-likelihood, and only completed the training until the likelihood increased less than (Δ = 0.01) in a single step. We used 50 initializations, each trained for an initial 30 steps.

Finally, we repeated this entire process a minimum number of 50 times and then continued until the solution with the highest likelihood was found at least 10 times in independent learning runs. This increased our confidence that we indeed had found a solution that could constitute a global maximum.

### 4.2. fMRI Datasets

In this project, we considered seven task-based and one resting-state fMRI datasets (see Table 1). The task-fMRI datasets refer to: (1) the *Multi-Domain Task Battery* (MDTB, King et al. (2019)); (2) a high-resolution version of the MDTB (*High-res MDTB* ; not yet published); (3) the *Nakai & Nishimoto* dataset (Nakai and Nishimoto, 2020); (4) a subset of the *Individual Brain Charting (IBC)* dataset (Pinho et al., 2018, 2020); (5) the *Shahshahni* dataset (Shahshahani et al., 2023); (6) the *Multi-Demand* dataset (Assem et al., 2022); and (7) the *Somatotopic* dataset (Saadon-Grosman et al., 2022). The first four datasets of the list include a broad range of task conditions from the perceptual, cognitive, motor, and social domains. In the first three datasets, tasks were randomly intermixed in each imaging session. In the *IBC* dataset, individual runs comprised only one task or a few tasks pertaining to a specific cognitive domain. The three last datasets of the list probe a more circumscribed array of functions: the *Shahshahni* dataset includes verbal working memory tasks (with forward and backward recall) and finger tapping tasks; the *Multi-Demand* dataset includes three executive function tasks (n-back, task-switch, a no-go); and the *Somatotopic* dataset probes foot, hand, glutes, and tongue movements. Lastly, as a resting-state fMRI dataset, we used the *Unrelated 100* subjects, which made publicly available in the *Human Connectome Project (HCP)* S1200 release (Van Essen et al., 2013).

The task-based datasets were pre-processed using either the *SPM12* software package (Wellcome Department of Imaging Neuroscience, London, UK) or the *FSL* library (Analysis Group, FMRIB, Oxford, UK). For every participant, an anatomical MRI image (T1- weighted MPRAGE, 1mm isotropic resolution) was acquired in one scanning session. FMRI data (time series acquired with Echo-Planar Imaging, T2*-weighted sequence using Blood-Oxygenation-Level-Dependent contrast) were realigned for head motion within each session, and for different head positions across sessions using the six-parameter rigid body transformation (Friston et al., 1995, Jenkinson et al., 2002). The mean functional image was then co-registered onto the anatomical image and this transformation was applied to all functional images (Ashburner and Friston, 1997, Greve and Fischl, 2009). No smoothing or group normalization was applied.

In parallel, the individual anatomical volumes were segmented into different tissue types (Ashburner and Friston, 2005), and the whole-brain plus gray-matter masks were derived from this segmentation. Each anatomical image was submitted to the standard recon-all pipeline from the *FreeSurfer* software (Fischl, 2012) to obtain a reconstruction of the individual cortical surfaces. Similarly, each anatomical image was processed using SUIT (Diedrichsen, 2006), which provided cerebellar segmentation and normalization. The cerebellar mask was derived from this segmentation and hand-corrected, whenever necessary, to ensure that voxels from occipital and inferior temporal cortices were not included.

A mass-univariate General Linear Model (GLM) was then fitted to the realigned functional data to estimate brain activation per imaging run. Each task condition was modeled as a boxcar function according to the onsets and duration of the given task condition. The corresponding boxcar function was then convolved with the canonical Hemodynamic Response Function (HRF) (Friston et al., 1998a,b). The whole-brain mask was applied to the realigned functional volumes to restrict the GLM to voxels inside the brain. Coefficients of the GLM were divided by the root-mean-square error (RMSE) for each voxel, resulting in individual volume-based maps of normalized activity estimates. These functional derivatives, obtained for each task condition and imaging run served as input to the fMRI dataset integration framework (see Section 4.3).

The resting-state data were pre-processed using the HCP minimal processing pipeline (Glasser et al., 2013), including structural registration, correction for spatial distortion, head motion, cortical surface mapping, and functional artifact removal (Smith et al., 2013, Glasser et al., 2013). For each imaging run, this resulted in 1200 time points of processed time series for each voxel of the standard MNI152 template (Van Essen et al., 2012) in the cerebellum. To generate the resting-state functional connectivity (rs-FC) fingerprint of the cerebellar voxels from the HCP data set, a group Independent Component Analysis (ICA) was applied. We first concatenated the preprocessed functional data temporally across subjects, sessions, and runs to create a single matrix. Then we used the group-ICA implemented in FSL’s MELODIC (Jenkinson et al., 2012) with automatic dimensionality estimation, resulting in 1072 group-level components. Sixty-nine signal components were identified from the first 300 ICA components as resting-state networks. Lastly, we regressed the 69 group network spatial maps into the subject-and-run-specific cortical time series, resulting in 69 cortical network time courses. The cerebellar rs-FC fingerprints were calculated as Pearson’s correlations of the cerebellar voxel time series with each cortical network time course.

### 4.3. Data structure and anatomical normalization

One important barrier to integrating task contrasts across different fMRI datasets is that these derivative measures are often stored in different atlas spaces (e.g. MNI, fsLR) and with different naming conventions, requiring specialized code for each dataset. To address this problem, we specified a data structure for fMRI derivatives using BIDS-derivative naming convention and file standards (Gorgolewski et al., 2016). For each dataset, we imported the task contrasts (estimates) for each subject, run, and condition that were estimated from minimally pre-processed, non-normalized, and un-smoothed, fMRI data (see Method 4.2). We then developed a toolbox that allowed the automatic and fast extraction of this data in any desired atlas space (surface- or volume-based), at any desired level of smoothing and aggregation across runs.

After extraction the resulting functional files are stored using the CIFTI format, resulting in fast and efficient loading times. For the current project, we extracted the cerebellar data in 3mm resolution, aligned to the MNI152NLin2009cSym template (Ciric et al., 2022), resulting in 5446 voxel locations in group space. The sampled functional data of all datasets were smoothed using a Gaussian kernel of 2mm standard deviation, except the *Somatotopic* dataset that used a 3mm smoothing kernel. The proposed file structure and code are available in a public repository (https://github.com/DiedrichsenLab/Functional_Fusion). The parcellations were visualized using a surface-based representation of the cerebellum (Diedrichsen and Zotow, 2015).

### 4.4. Synthetic datasets for simulation

To validate the proposed framework, we ran several simulations (Results 2.2, 2.3) on synthetic datasets. To generate individual brain organization maps (**U***^s^*), we used a Markov random field of rectangular 50 × 50 grid with a 4-neighbor connectivity scheme. Each grid point represented a brain location and could take one of *K* values (a.k.a Potts Model (Wu, 1982)). We first generated an artificial smooth group probability map (Fig. S2a) by selecting *K* centroids *µ_k_* at random locations, and assigning the bias parameters of the spatial arrangement model *η_i,k_* for the node a location *x_i_*to be:

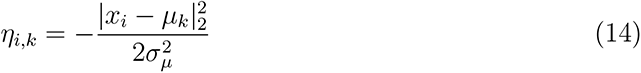

where 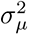 controls the smoothness of the group map (see Supplementary Fig. S2b).

The individual maps **U***^s^*were then sampled from the Potts model where the local probability *ψ_i,j_* between two vertices *i* and *j* was set to

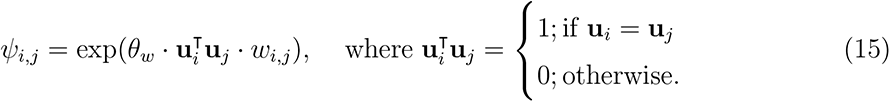

The pairwise weight of two vertices *w_i,j_* (*w_i,j_* = *w_j,i_*) indicates whether *i* and *j* are neighbouring vertices (*w_i,j_*= 1 if *i* and *j* are neighbours; *w_i,j_*= 0 otherwise). The temperature parameter *θ_w_* controls how strong the spatial co-dependence between neighbouring vertices is. A higher *θ_w_* encourages that the two neighbouring nodes are more likely to be assigned to the same parcel, enforcing the overall local smoothness of the map (Supplementary Fig. S2c). Ultimately, the individual maps were generated using vertex-wise Gibbs sampling after a burn-in of 20 iterations across all vertices.

We then generated synthetic functional data **Y***^s^* for each participant based on their individual parcellation maps. Rather than using a von Mises-Fisher distribution, we wanted to generate data that had both an amplitude and direction. Additionally to the region-specific mean direction of the response *v_k_*, we therefore introduced a non-negative region-specific signal strength parameter, *λ_k_*. The data for each voxel was generated from:

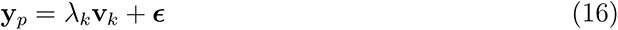

where ***ɛ*** was a normal random vector with variance 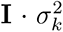. These parameters allowed us to control the signal and noise levels in each region separately. After normalization of the data to unit length, the generated data conformed approximately to a von Mises-Fisher distribution with mean **v***_k_* and concentration 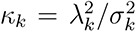. Ultimately, a synthetic dataset consisting of *N* task observations was generated for *P* brain locations and *S* subjects.

For the simulation in Results 4.2 and 4.3, the bias terms for the Potts model were generated with 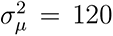, while the true number of parcels in the group map and fitting model were both set to *K* = 20. Then, we sampled 10 individual maps **U***^s^* from the group map with local connection weights *w_i,j_* = 1.5. These individual maps are further used to sample synthetic data from two sessions **Y***^s,^*^1^ (session 1, *N* = 40 tasks), **Y***^s,^*^2^ (session 2, *N* = 20 tasks) and a test set **Y***^s^_test_* (*N* = 120 tasks) with equal signal strength *λ_k_* = 1.1 for all functional regions. The *λ_k_* might be changed depending on specific simulations (see Results 4.2 and 4.3).

### 4.5. Evaluation measures for probabilistic atlas

The distance-controlled boundary coefficient (DCBC, Zhi et al. (2022)) is an unbiased evaluation criterion for brain parcellation, which allows the direct comparison of brain maps generated from different modalities (i.e resting-state, task-based, and anatomical) and different number of parcels. The coefficient controls for the intrinsic smoothness of brain data, which is biased in other evaluation metrics (Gordon et al., 2016, Rousseeuw, 1987). The DCBC method solves this problem by binning all vertex pairs based on their spatial distance and only comparing the Pearson’s correlation for within-parcel pairs and between-parcel pairs for the same distance. Then, the DCBC value is calculated as the average correlation differences, weighted by reliability across distances. The spatial distance is calculated as the Euclidean distance between the center of each voxel pair in the atlas volume space. The underlying functional profiles for calculating the correlations of voxel pairs are the associated betas weights in a task-based dataset. A higher DCBC value of a parcellation indicates that this parcellation predicts the functional boundaries well on the tasks of the dataset being used.

### 4.6. Computational setup

Model training and evaluations were performed on either an NVIDIA 1080Ti GPU with Python 3, CUDA 11.3, and PyTorch 1.10.2 or on NVIDIA GRID A100-10C GPU with Python 3, CUDA 11.6, and PyTorch 1.13.1. For the fMRI datasets, all data were preprocessed and extracted on an Intel i7-8700 CPU with NumPy 1.24.0, NiBabel 4.0.2, neuroimagingtools 0.5.0. Other detailed requirements and parameters used for the data processing pipeline are available in the respective repositories (see Code availability).

## 5. Data availability

The raw data for the fMRI studies used in this project are publicly available on https://openneuro.org/ for the studies listed in Table 1.

## 6. Code availability

The code for the hierarchical Bayesian parcellation framework is publicly available as the GitHub repository https://github.com/DiedrichsenLab/HierarchBayesParcel. The organization, file system, and code for managing the diverse set of datasets is available in a separate repository https://github.com/DiedrichsenLab/Functional_Fusion. The paper-specific code for generating the functional probabilistic parcellations for the cerebellum, as well as running the simulation presented in this paper is available at https://github.com/DiedrichsenLab/FusionModel.

## 7. Acknowledgements

This study was supported by a Discovery Grant from the Natural Sciences and Engineering Research Council of Canada (NSERC, RGPIN-2016-04890), and a project grant from the Canadian Institutes of Health Research (CIHR, PJT 159520), both to J.D. Additional funding came from the Canada First Research Excellence Fund (BrainsCAN) to Western University. Special thanks to N. Grosman, R. Buckner, M. Assem, and J. Duncan for sharing their datasets before the official public release.

## 8. Author contributions

All authors were involved in conceptualizing the framework and writing of the paper. D.Z. and J.D. developed the computational details. D.Z., A.L.P., L.S., C.N., and J.D. oversaw the pre-processing of various datasets.

## Supplementary Materials and Figures

### Parameter estimation of full model

In this section, we provide details of model parameter estimation for the full EM algorithm. The complete expected log-likelihood Σ*_s_*⟨log *p*(**Y***^s^,* **U***^s^*; ***θ***)⟩*_q_* can be decomposed into expected emission log-likelihood L*_E_* and expected arrangement log-likelihood L*_A_*, where ⟨·⟩*_q_* denotes the expectation with respect to distribution *q*. Similarly, the model parameter *θ* can be subdivided into *θ_E_* and *θ_A_* and can be estimated within their corresponding models (Methods 4.1). This unique model structure yields the following learning EM process:

#### Emission model E step

Suppose for a single dataset **Y***^n^* is a *S* × *N* × *P* tensor for *S* subjects (*S* is the number of subjects in S) of *N* data observations across *P* voxels. The brain activation of a voxel for a single subject **y*_i_****^s^* is a *N* -long vector. If the task design has repeated measurements of the same *M* conditions (e.g. in a single imaging run), the user can specify this over a *N* × *M* design matrix *X* (*M* is the number of unique task conditions). To account for the situation that **y*_i_****^s^* consists of multiple partitions, which could be imaging sessions or runs, we used an *N* -dimensional partition vector to divide *N* observation into *J* independent partitions. Therefore, if we combine the data across repeated measurements and different partitions, the resultant data **ỹ*_i_****^s^* would be a sum of normalized data in each partition *j* as,

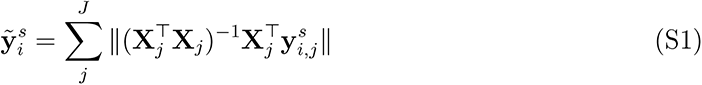

However, we can also treat the different repetitions as independent observations, meaning that the resultant data is normalized to length 1 across *J* independent partitions. This is also the case with the Type 1 model, in which the imaging sessions are simply concatenated. Hence, the expected emission likelihood L*_E_* of a mixture of *k*-classes vMF distribution in eq. 11 is modified and updated at (*t* + 1) iteration by:

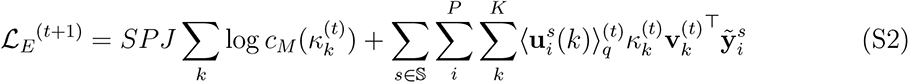

As a sufficient statistic, it should be noticed that the resultant summed vectors **ỹ**_i_*^s^* become a *M* -dimensional vector but its magnitude is not 1 anymore. Therefore, the normalizing constant will be computed in *M* -dimensional correspondingly, denoted as log *c_M_* (*κ_k_*).

#### Arrangement model E step

Expanding eq. 10 and 7, the expected posterior under the proposal distribution *q* at (*t* + 1) iteration are updated as,

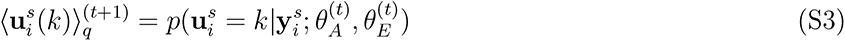

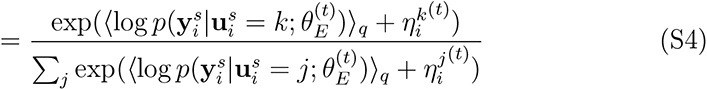

where *η^k^* is defined in Methods.

#### Arrangement model M step

Expanding the expected arrangement log-likelihood in eq.10, we obtain the derivatives with respect to the parameters *θ_A_* := {*η_i,k_*}:

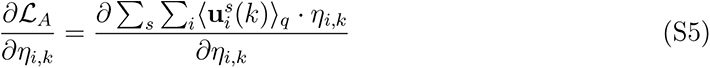

By setting this derivative to zero, we can obtain the following parameter updates:

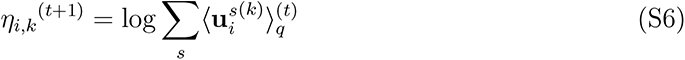

#### Emission model M step

To update the parameters *θ_E_* of the vMF mixture in the M-step, we need to maximize L*_E_* in respect to the parameters in vMF mixture *θ_k_* = {**v***_k_, κ_k_*}. First, we update the mean direction **v***_k_*, where we get the intuitive update :

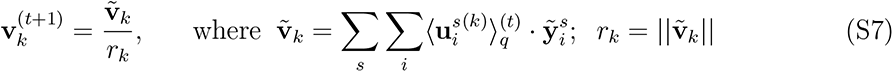

The updates of the concentration parameters *κ_k_* are more difficult in particular for high dimensional problems, since it involves the inverting ratio of two Bessel functions. Therefore, we here use an approximate solution suggested by Banerjee et al. (2005) and Hornik and Grün (2014). In our specific case, we want to integrate the evidence across *s* = 1*, …, S* subjects and *i* = 1*, …, P* voxels, with each subject and voxel may have *J_i_^s^* partitions. Under this assumption, we can **(1)** learn a common *κ* across classes by restricting *κ_k_* to be equal, as:

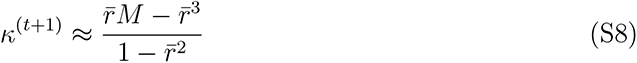

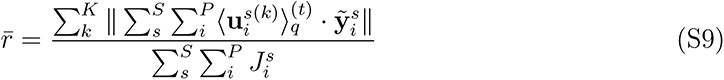

which is used in Type 1 and Type 2 model learning.

Alternatively, we can **(2)** learn *k*-class specific kappa *κ_k_* by relaxing the constraint as:

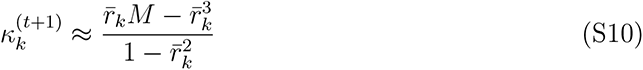

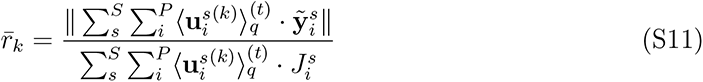

which will be used as the parameter estimates for the Type 3 regions-specific emission model.

**Figure S1:**
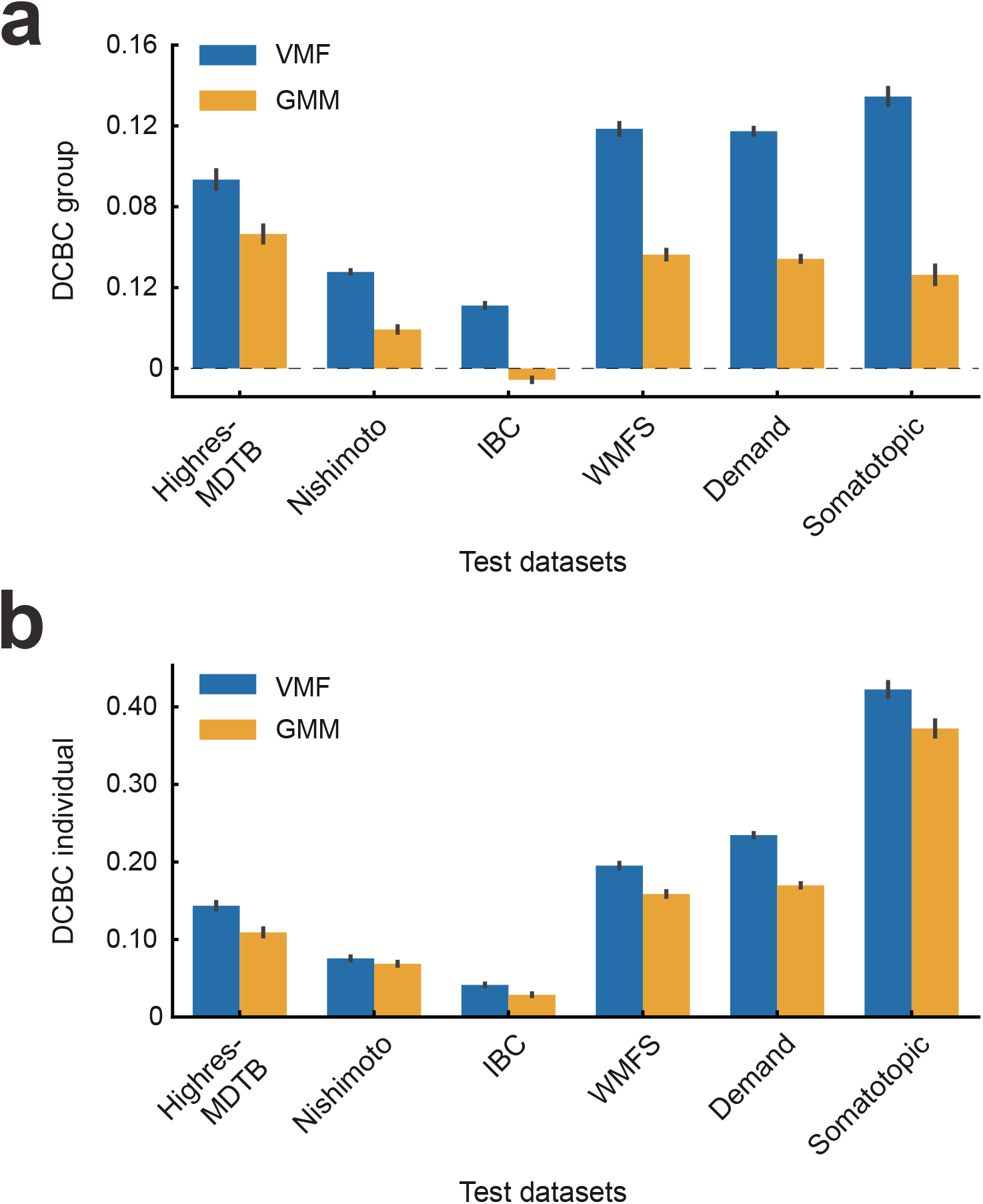
Comparison of the performance between the parcellations derived from Gaussian Mixture Model (GMM) and von Mises-Fisher Mixture model (VMF). **(a)** The averaged DCBC value of the group parcellation maps trained by GMM or VMF mixture model across subjects in the test dataset. **(b)** The averaged DCBC value of the individual parcellation maps trained by GMM or VMF mixture model across subjects in the test dataset.

**Figure S2:**
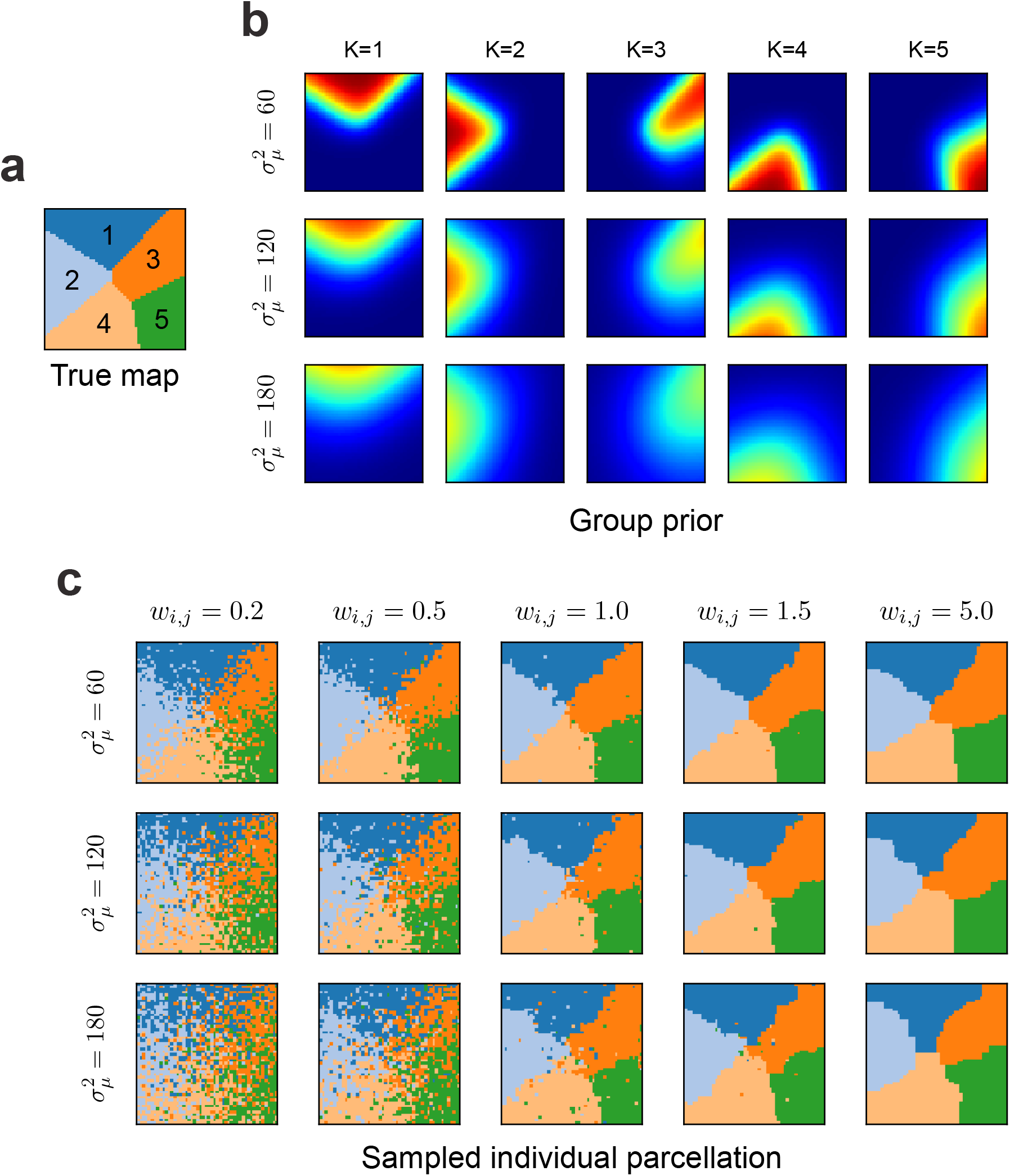
The synthetic dataset. **(a)** The random true group map with 5 parcels. **(b)** The group prior controlled by smoothing kernel at different levels for all 5 classes. **(c)** The example individual parcellation maps generated by different parameters

**Figure S3:**
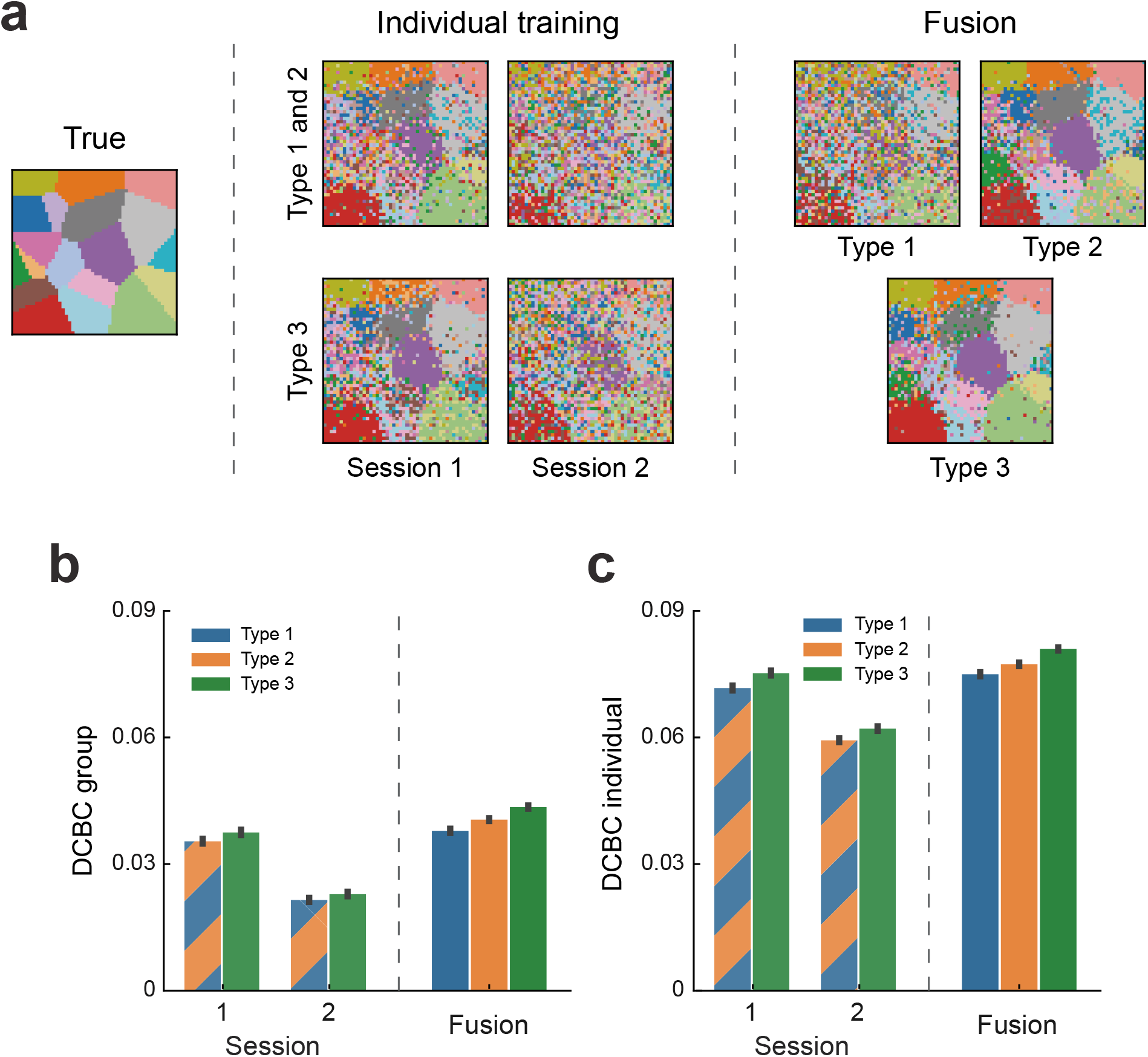
Simulation on two synthetic sessions fusion with similar task activation using Type 1, 2, and 3 emission models. **(a)** The comparison of model reconstruction performance of group parcellations learned on synthetic session 1 or 2 standalone vs. the ones learned fusion using type 1, 2, or 3 models. **(b)** The mean DCBC value of the group map learned from session 1 or 2 only or learned by fusion using type 1, 2, or 3 models. **(c)** The mean DCBC value of individual maps across all participants when learned from session 1 or 2 only or learned by fusion using type 1, 2, or 3 models. Error bars indicate SEM across 100 times simulation.

**Figure S4:**
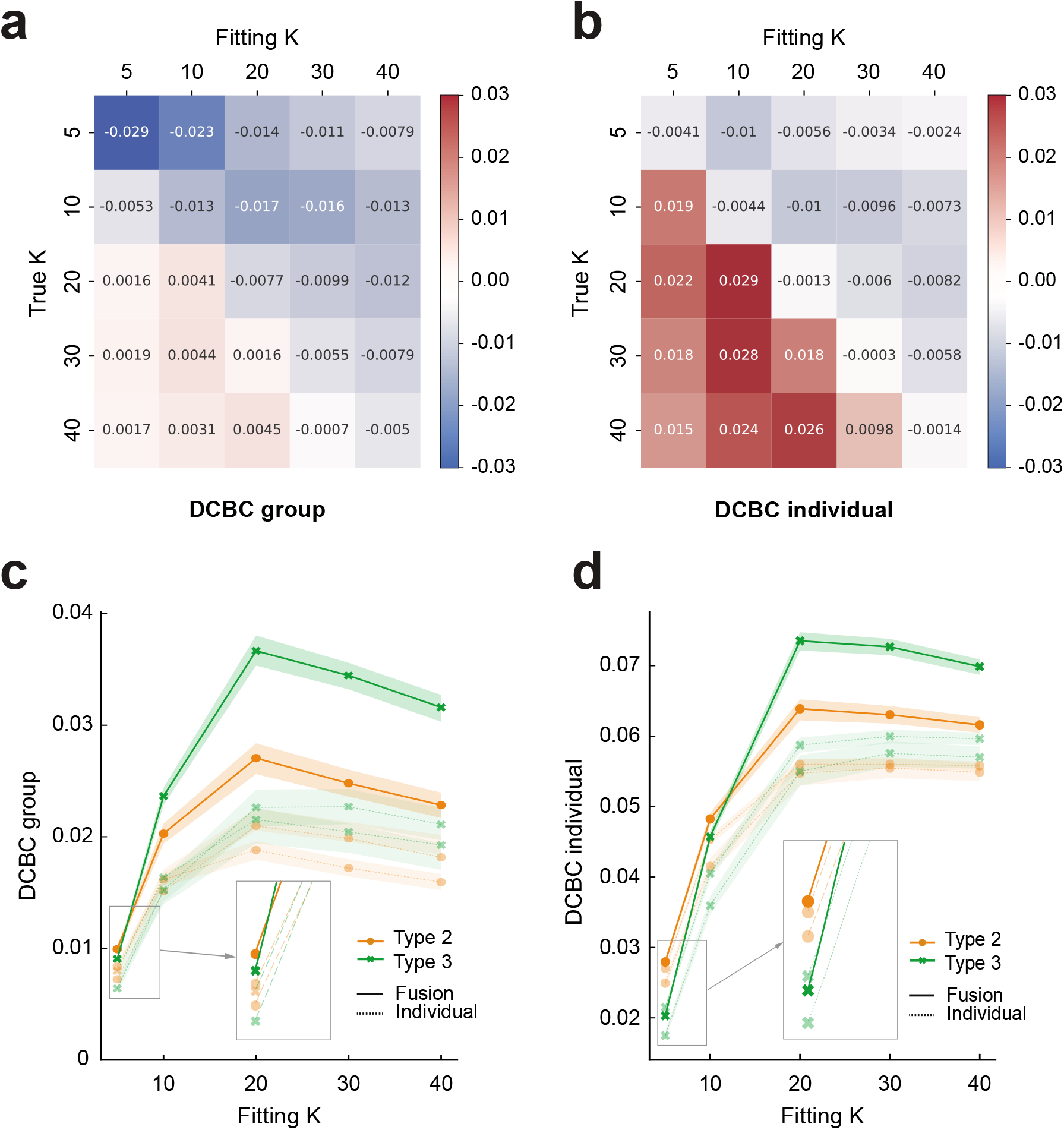
Comparing the performance of Type 2 and 3 models when the number of parcels *K* used for fitting is different from the true *K* in the simulation. (a) The difference of the mean DCBC value between the group map trained on a synthetic dataset using Type 2 and Type 3 models with different fitting *K* and ground true *K*, which tested on an independent synthetic test set. A positive value on the grid indicates the Type 2 model outperforms the Type 3 model, while negative values mean the opposite. **(b)** The difference of the mean DCBC value between the individual maps. **(c)** The mean DCBC value for the group map learned from individual synthetic datasets only or learned by fusion using the Type 2 or 3 models for *K* = 5 to *K* = 40 when the true *K* = 20. The error shade indicates the standard error across 100 simulations. **(d)** The mean DCBC value for the individual maps.

**Figure S5:**
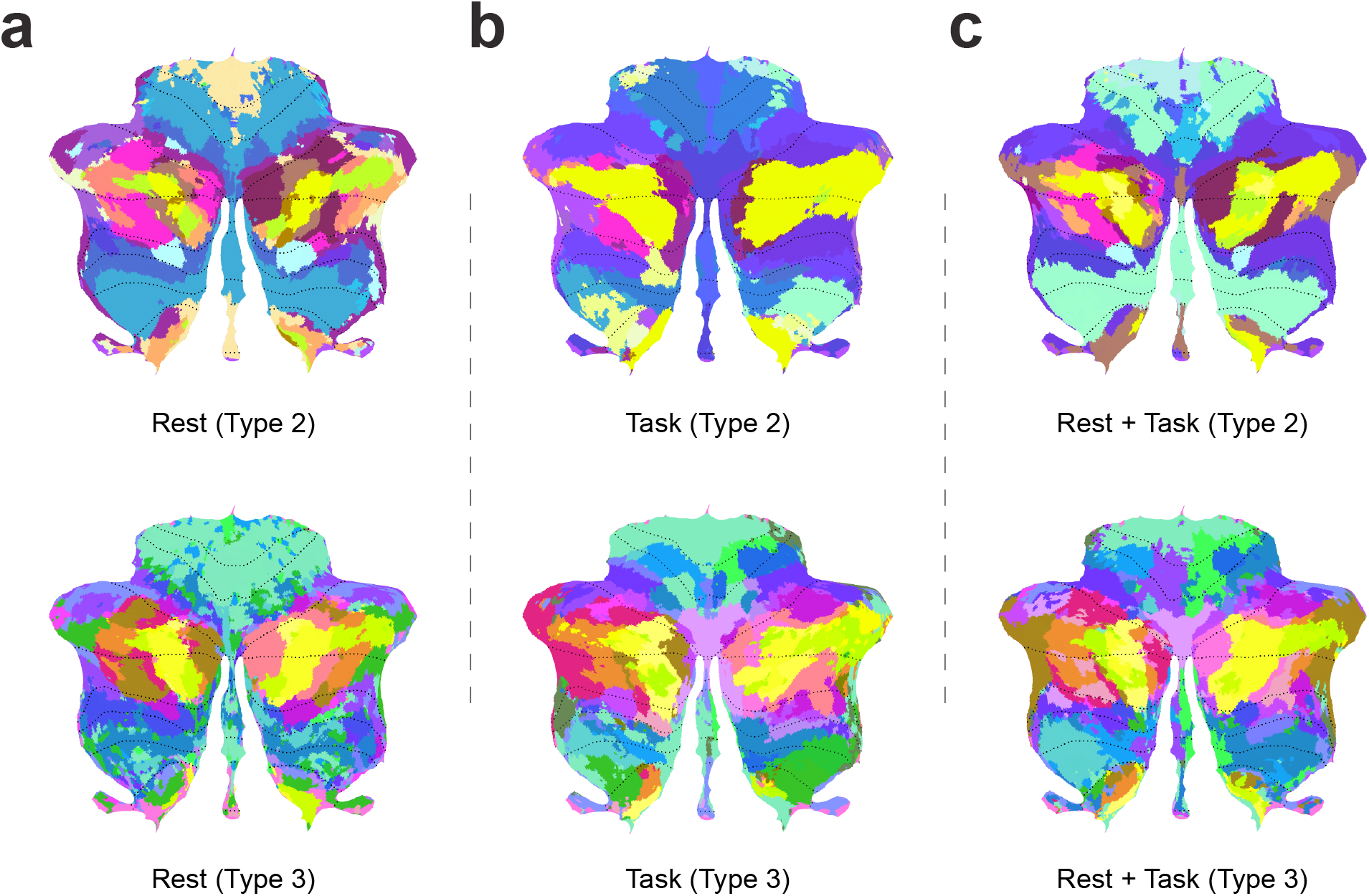
The visualization of the learned group maps (*K* = 34). **(a)** The maps were trained on a pure resting-state dataset *HCP-Unrelated 100* using the Type 2 or 3 fusion model. **(b)** The maps were purely trained on task-based datasets using the Type 2 or 3 fusion model. The task datasets are *MDTB*, *Highres-MDTB*, *Nakai&Nishimoto*, *IBC*, *WMFS*, *Demand*, *Somatotopic*. **(c)** The maps were trained on the combination of resting-state and all task-based datasets. The colors for the parcels are aligned in each type of model, where two similar colors in RGB space indicate the two parcels have similar task activation responses on average.

